# Transcriptomic Profiling of High vs. Low Flow Regions of Mouse and Human Trabecular Meshwork

**DOI:** 10.64898/2026.07.23.740307

**Authors:** Cydney A. Wong, A. Thomas Read, Micah Chrenek, Guorong Li, W. Daniel Stamer, Levi B. Wood, C. Ross Ethier

**Affiliations:** Wallace H. Coulter Department of Biomedical Engineering, Georgia Institute of Technology and Emory University, Atlanta, Georgia, USA; Department of Ophthalmology, Emory University, Atlanta, Georgia, USA; Department of Ophthalmology, Duke University, Durham, North Carolina, USA; George W. Woodruff School of Mechanical Engineering, Georgia Institute of Technology, Atlanta, Georgia, USA

**Keywords:** Trabecular Meshwork, Segmental Outflow, Gene expression, Transcriptomics, Glaucoma

## Abstract

**Purpose:** Aqueous humor outflow through the trabecular meshwork (TM) is segmental, demonstrating high flow (HF) and low flow (LF) regions. Here, we investigate transcriptomic differences between flow regions in naïve mouse and non-glaucomatous human TM tissue to better understand intraocular pressure (IOP) regulation.

**Methods:** Human eyes (<6 hours postmortem) from two donors were perfused with a fluorescent tracer to identify HF and LF regions and fixed. In parallel, one pair of 8-month-old C57Bl/6J mouse eyes were similarly processed. Sagittal sections from HF and LF regions underwent whole-transcriptome spatial profiling. Differential expression and gene set variation analysis were conducted to identify transcript and pathway-level differences between HF vs. LF TM. Selected targets were validated by immunolabeling.

**Results:** Genes with relevance to TM outflow were identified as significantly differentially expressed in the human dataset, including ADAM metallopeptidase domain 15 (*ADAM15*), vimentin (*VIM*), chitinase 3-like 1 (*CHI3L1*), and several transcription factors (e.g., *FOS, JUNB, ESR1*). Pathways related to epigenetic modifications were also differentially enriched in human eyes. In both human and mouse eyes, myocilin was significantly upregulated in HF regions, despite greater protein labeling in LF regions of human eyes. In both species, rho-kinase signaling pathways showed increased enrichment in LF regions, while cell stress pathways, and TNF-α signaling were increased in HF TM.

**Conclusions:** HF regions maintain a more active stress response that facilitates greater outflow, whereas LF regions exhibit more matrix accumulation and contractility. This characterization of segmental flow regions can inform future studies to target trabecular outflow and lower IOP.

## 1. Introduction

Primary open-angle glaucoma (POAG) is the leading cause of irreversible blindness, and the main risk factor for developing POAG is elevated intraocular pressure (IOP) due to increased resistance to aqueous humor (AH) outflow in the trabecular meshwork (TM) and inner wall of Schlemm’s canal (SC). Aqueous humor outflow is non-uniform around the circumference of the eye, i.e. there are high flow (HF) and low flow (LF) regions of the TM. This phenomenon, known as segmental outflow, has been extensively demonstrated using methods that involve introducing a visible tracer into the anterior chamber. The tracer follows the natural flow of AH through the outflow tract, with greater amounts of the tracer accumulating in high flow regions and less tracer in low flow regions. Previous studies have demonstrated segmental variability in outflow using tracers as varied as cationic and anionic ferritin^1–3^, quantum dots^4–6^, fluorescent nanospheres and microspheres^7–10^, and pigment^11^; fluorescent lentiviral reporters^12^; and fluorescein angiography^13–15^. While tracer size and surface chemistry may influence tracer location within the TM layers, localization to high flow regions around the circumference seems to be independent of these factors^5^.

Segmental flow has been observed in both normal and glaucomatous human eyes^1,9,16,17^, as well as porcine eyes^13,14^ and a variety of mouse strains^6,10,18^. While some association of increased outflow near collector channels has been reported^11^, this cannot fully explain the outflow variation observed because the collector channels are fairly evenly distributed around the limbus in human eyes, and segmental flow patterns are dynamic and can change with time^10^ and pressure elevation^9^.

While the mechanisms governing segmental outflow are not fully understood, previous work has identified both molecular and biomechanical differences in TM cells from high vs. low flow regions. For example, expression levels of extracellular matrix (ECM)-related genes, including versican, SPARC, collagens, and MMPs, differ between segmental flow regions^4,5,19^. More recently, transcriptomic profiling in high vs. low flow regions of human donor eyes revealed ADAM15 as a major differentially expressed gene, along with several other matrix and cell-adhesion related genes^20^. Further, cells extracted from HF and LF regions of human TM demonstrate differences in stiffness, matrix deposition, and gene expression^21–23^ in vitro. In mice, the molecular signatures are not as well understood; however, mice are excellent models for understanding changes in segmental outflow in a living system^10^, as well as the effects of genetic knockouts^19^ and potential drug treatments^24^. Thus, an understanding of segmental outflow at the transcriptomic and protein levels in both human and mouse eyes remains a major knowledge gap. Deeper understanding of differences in gene expression, signaling pathways, and protein levels between segmental flow regions could provide novel insights into the regulation of aqueous humor outflow and glaucoma pathophysiology.

In this study, we used spatial profiling to identify transcriptomic differences between high and low flow regions of the mouse and human TM and further investigated differences at the protein level by immunofluorescence staining in high vs. low flow regions from human TM. This allowed us to identify similarities and differences between mouse and human segmental flow biology and further our understanding of the mechanisms of segmental outflow.

## 2. Methods

### 2.1. Human Donor Tissue

Human donor eyes were obtained from Lions World Vision Institute in Tampa, FL. Both eyes from one donor (72-year-old male) and one eye from a second donor (73-year-old female) were received and processed within 5 hours postmortem (**Table 1**). No ocular abnormalities were observed, and neither donor had a history of eye disease. The human tissue experiments complied with the guidelines of the ARVO Best Practices for Using Human Eye Tissue in Research.

**Table 1:** Human Donor Tissue Samples. Neither donor had a history of ocular disease or diabetes. Processing of the samples began immediately upon receipt.

| Donor | Eyes analyzed | Age, Sex | Race | Postmortem enucleation time | Time received post enucleation |
| --- | --- | --- | --- | --- | --- |
| 1 | OD, OS | 72y, M | White | 4 hours | 40 minutes |
| 2 | OS | 73y, F | Unknown | 4 hours | 1 hour |

### 2.2. Human Tracer Perfusion and Tissue Processing

After removing connective tissue, fat, muscle, and conjunctiva, each eye was secured with the limbus exposed in a moist environment with phosphate buffer saline (PBS) at room temperature. The eye was cannulated with two needles: one connected to an input reservoir, and another connected to an output reservoir. One needle was advanced through the cornea parallel to the corneal surface until its tip resided in the anterior chamber, while the second needle was inserted through the cornea midway to the limbus so that the tip of the needle rested just beneath the iris into the posterior chamber. The anterior chamber was then exchanged with approximately 2 mL of Dulbecco’s phosphate buffered saline with added 5.5 mM glucose (DBG), and then the eye was perfused with DBG for 10 minutes at approximately 18 mmHg. The anterior chamber contents were then exchanged with a solution of fluorescent, carboxylate-modified 100 nm Fluospheres® (515 nm emission, Molecular Probes, Eugene, OR, USA) diluted 1:5,000 in 5 mL degassed DBG, followed by a 15-minute perfusion at approximately 18 mmHg. Finally, the anterior chamber was exchanged with 4% paraformaldehyde in PBS, followed by a 15-minute perfusion at approximately 18 mmHg to fix outflow tissues. The eye was dissected into quadrants, the ciliary body, choroid, and iris were resected, and the quadrants stored in RNALater (Thermofisher) at 4 °C.

Tissue quadrants were washed in PBS, further dissected into wedges, resected to expose the TM, and imaged on a Leica DM6 microscope with a GFP filter cube to identify regions with the highest fluorescent signal (HF) and lowest fluorescent signal (LF) based on tracer distribution around the limbus. Once target HF and LF regions were identified and annotated using the microscope’s software, wedges were cryoprotected, embedded in optimal cutting temperature compound (OCT), snap frozen in 2-methylbutane cooled with liquid nitrogen, and stored at −80 °C.

### 2.3. Human Whole Transcriptome Digital Spatial Profiling

Sagittal 8 µm thick cryosections were collected from target HF and LF regions of the anterior chamber and arranged on a SuperFrost Gold Plus slide. Each slide contained at least 12 HF and LF sections from one eye (**Figure 1B**). Slides were stored at −80°C and shipped to Bruker Spatial Biology Technology Access Program (TAP) in Seattle, WA for spatial profiling with the GeoMx Digital Spatial Profiler (DSP).

**Figure 1:**
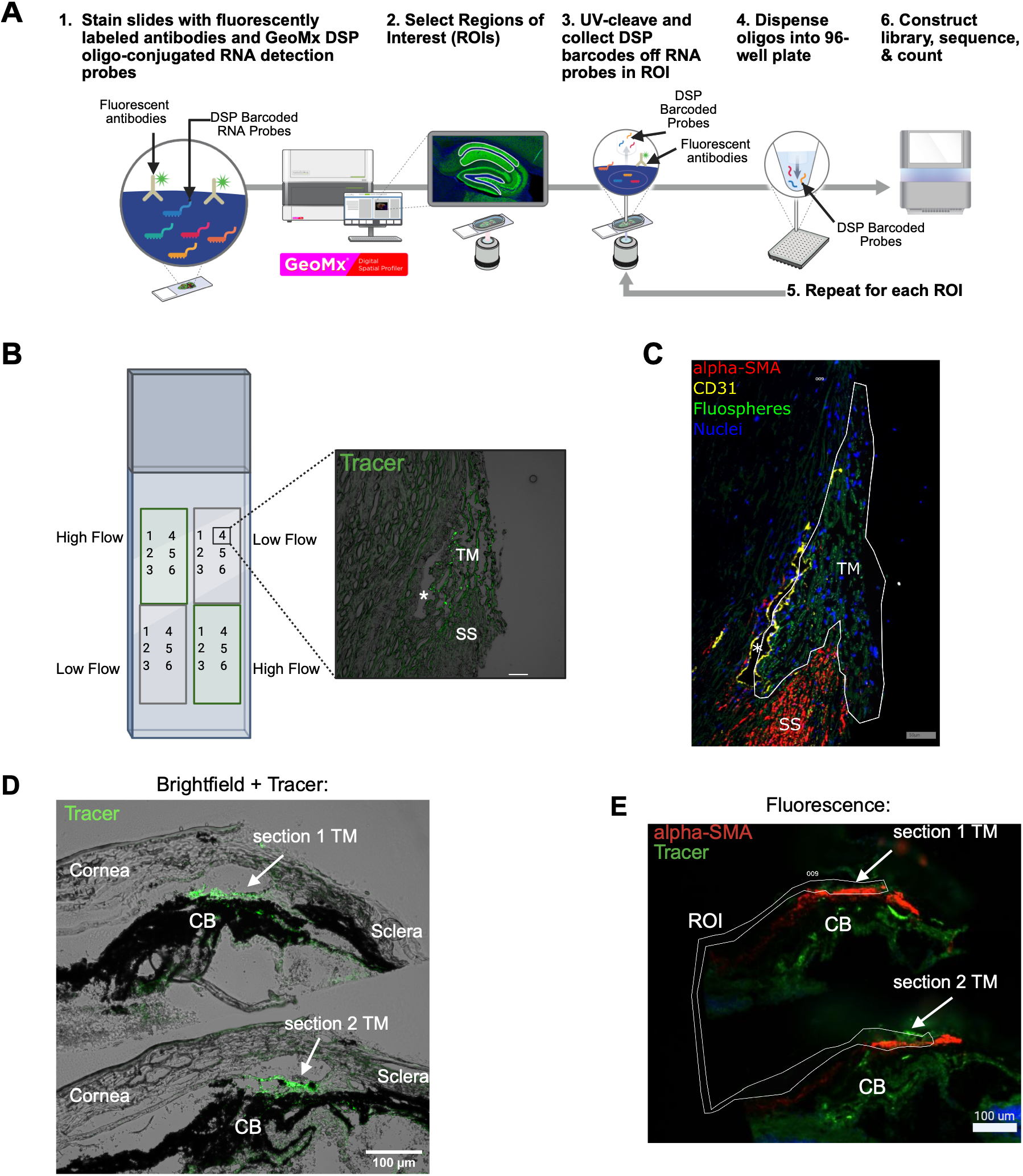
GeoMx Spatial Profiling in Human and Mouse TM.^25^ **(A)** Overview of GeoMx DSP workflow. Slides are incubated with the DSP Whole Transcriptome Atlas RNA Probes, and tissues are labeled with fluorescent antibodies to provide tissue landmarks. When the slides are in the instrument, a user can manually select ROIs based on the landmarks. Barcodes are UV-cleaved from each ROI, collected into a 96-well plate, and sequenced using an Illumina NovaSeq. (Figure is modified from Bruker Spatial Biology). **(B)** Sample slide layout containing 12 HF and 12 LF sections from a single human eye, with the callout box exhibiting an example section from one eye. Green flow tracer is visible in the TM, and SC lumen is indicated by an asterisk (*). SS = Scleral spur, scale bar = 50 µm. **(C)** Representative ROI selection on human tissue section labeled with morphology markers. The white outline indicates the ROI, and the labels for a-SMA, CD31, and nuclei are shown, in addition to the flow tracer (Fluospheres) in green. Scale bar = 50 µm. **(D)** A representative pair of mouse anterior segment tissue sections, shown as an overlay of the brightfield and fluorescence channel to visualize tracer location in the TM. After imaging sections, most of the cornea and iris were scraped away with a razor blade so that the ROI could be drawn connecting the TM from both sections. **(E)** Fluorescent image of sections in panel D, labeled with α-SMA (in red) and CD31 (not shown). The ROI, outlined in white, includes the TM from both sections. CB = ciliary body.

Slides were processed by the TAP team using standard protocols. In short, the sections were dried and baked at 37 °C for one hour, then incubated in 4% paraformaldehyde for 60 minutes at room temperature. Slides were then washed and dried before automated target retrieval with 0.1 ug/mL proteinase-K for 15 minutes and epitope retrieval solution at 100 °C for 10 minutes. Finally, slides were hybridized overnight with the GeoMx Human Whole Transcriptome Atlas (WTA) nucleic acid probes coupled with photocleavable oligonucleotide tags (**Figure 1**).

The slides were also stained with fluorescent antibodies against α-SMA (Abcam ab202368) and CD31 (R&D Systems AF3628), and nuclei were labeled with DAPI (Thermofisher). The fluorescent tracer labeling segmental flow regions was also visible in the green (GFP) channel and was used to help identify the TM. Once loaded in the DSP instrument, we remotely accessed the system and viewed fluorescent labeling on the slides to identify the TM and hence draw regions of interest encompassing the TM. Specifically, the CD31 labeling of SC endothelial cells, α-SMA labeling of scleral spur and ciliary muscle, and green flow tracer in the TM were used to guide ROI selection (**Figure 1C**). Then, UV light was delivered only to the outlined regions of interest (ROI), and the barcoded RNA oligos in the WTA library were photocleaved. Barcodes for each ROI were then collected into a 96-well plate and sequenced on an Illumina NextSeq 2000. FASTQ files generated from sequencing were processed with the Nanostring GeoMx pipeline to obtain transcript count data for each ROI.

### 2.4. Human Transcriptomic Data Analysis

A limit of quantification (LOQ) was determined for each ROI based on counts from negative probes included in the atlas. To remove low-expressed genes from our data, any genes not detected above the LOQ in 5% of the ROIs were removed from analysis. After filtering, we were left with 8,299 genes remaining from the over 18,000 genes included in the whole transcriptome atlas. Q3 normalization was performed in the GeoMx data analysis suite to combine count data from all samples, followed by variance-stabilizing normalization (VSN) in RStudio (“vsn” package, v. 3.64.0). Principal component analysis (PCA) was performed to reduce the dimensionality of normalized count matrix (8,299 genes x 72 ROIs) using the “stats” package (v. 4.2.0) in RStudio. Given the high dimensionality of the dataset and the expectation of subtle variation among samples derived from the same tissue type, we retained only the first five principal components for visualization and calculation of percent variance captured.

Differential expression analysis was performed using the limma package in R (limma v. 3.52.4). A linear model, with flow region as a fixed effect, was used to find differences between high and low flow regions, where each of the ROIs were treated as individual samples and donor effects were modeled as random using the “duplicateCorrelation” function to account for repeated measures from the same donor. Empirical Bayes moderation was used to compute statistics and adjusted p-values were calculated using the Benjamini-Hochberg (FDR) method. However, no genes met the FDR threshold for significance (FDR < 0.05), so genes with unadjusted p < 0.05 are reported as significantly differentially expressed to facilitate interpretation of expression trends in this small dataset of three donor eyes. PCA was also performed on the donor-corrected normalized count data to confirm successful removal of donor effects. Hierarchical clustering of differentially expressed genes was performed using “ComplexHeatmap” (v. 2.12.1).

To assess the representation of genes with known association to IOP elevation in our dataset, we compiled a list of candidate genes from one genome-wide analysis that identified loci associated with POAG^26^ and two genome-wide analyses that identified loci significantly associated with IOP elevation^27,28^. Genes associated with each locus were combined into a non-redundant list and cross-referenced with the normalized expression data and differential expression analysis results to determine which IOP-associated genes were detected and differentially expressed between HF vs. LF regions.

Gene set variation analysis (GSVA) was used for pathway-level analysis^29^ using the Molecular Signatures Database (MSigDB) C2 collection of curated gene sets (v7.5.1)^30^. Statistical differences in enrichment scores for each gene set between subject groups were computed by comparing the true differences in means against a null distribution obtained by permuting the gene labels and re-computing the GSVA 1,000 times. Statistical comparisons were based on gene expression counts that had been adjusted to account for repeated measures from each donor, so each ROI was treated as an individual sample.

### 2.5. Mouse Tissue Preparation and GeoMx Spatial Profiling

All animal procedures were approved by the Institutional Animal Care and Use Committee (IACUC) of Duke University and were consistent with the ARVO Statement for the Use of Animals in Ophthalmic and Vision Research. An 8-month-old female C57BL/6J wild-type mouse purchased from the Jackson Laboratory (Bar Harbor, Maine, USA) was perfused bilaterally in vivo with a 1:15,000 dilution of fluorescent, carboxylate-modified 100 nm Fluospheres® (515 nm emission, Molecular Probes, Eugene, OR, USA) in PBS at a constant pressure of 15 mmHg for 40 minutes. Both eyes were fixed by immersion in 4% paraformaldehyde (Thermofisher) for one hour before being transferred into Diethyl pyrocarbonate (DEPC)-treated PBS.

Downstream tissue processing and preparation for GeoMx profiling was performed similarly to the human samples, with adjustments made to tissue layout on the slide to capture cells from two sections in each ROI to obtain optimal sensitivity (**Figure 1D, E**). Details on mouse sample preparation can be found in the Supplementary Files.

### 2.6. Mouse Transcriptomic Data Analysis

A limit of quantification (LOQ) was determined for each ROI based on counts from negative probes included in the mouse whole transcriptome atlas. To remove low-expressed genes from our data, any genes not detected above the LOQ in 20% of the ROIs were removed from analysis. After filtering, we were left with 3,338 genes remaining from the over 20,000 genes included in the mouse whole transcriptome atlas. Q3 normalization was performed in the GeoMx data analysis suite. In RStudio, principal component analysis (PCA) was performed to reduce the dimensionality of normalized count data and hierarchical clustering and heatmap generation was performed using the “ComplexHeatmap” package (v. 2.12.1).

Differentially expressed genes were identified in the GeoMx Data Analysis Suite using a t-test with FDR-adjusted p-values; however, due to the small sample size, none of the genes were significant after FDR-adjustment, so raw p-values are shown on all plots, and genes with unadjusted p < 0.05 were considered differentially expressed. Gene set variation analysis (GSVA) was used for pathway-level analysis ^29^ using the Molecular Signatures Database (MSigDB) C2 collection of curated gene sets ^30^. Statistical differences in enrichment scores for each gene set between subject groups were computed by comparing the true differences in means against a null distribution obtained by permuting the gene labels and re-computing the GSVA 1,000 times, and these values were FDR-adjusted.

### 2.7. Immunofluorescence Microscopy

Sagittal 10 µm thick anterior segment sections from HF and LF regions of each human donor eye were collected with a CryoStar NX70 cryostat (Thermofisher). Prior to staining, all slides were baked at 37 °C for one hour.

Sections to be labeled with antibodies against myocilin, fibronectin, SPARC, collagen I, or collagen VI were permeabilized with 0.2% Triton X-100 and blocked in 10% goat serum for 30 minutes at room temperature. Sections were then incubated overnight at 4 °C with primary antibodies (see **Supplemental Table 1**), followed by overnight incubation at 4 °C with either Alexa Fluor 546 goat anti-rabbit secondary antibody (Invitrogen, A-21245; 1:200 dilution) or Alexa Fluor 546 goat anti-mouse secondary antibody (Invitrogen, A-11030; 1:200 dilution), depending on the primary antibody. Each slide also included 1-2 negative control sections prepared in the same way, except that the primary antibody was omitted. Nuclei were counterstained using NucBlue FixedCell Stain ReadyProbes (Invitrogen, Waltham, MA, USA) according to the manufacturer’s instructions. Glass coverslips were mounted with Prolong Gold Antifade mounting medium (Invitrogen, Waltham, MA, USA). Images were acquired on a Leica DM6 epifluorescence microscope with a 40× objective lens to obtain 6.2 pixel/µm resolution extended depth of field images from a z-stack. Identical exposure times and gain settings were used for all samples. Fluorescence signals were detected using the GFP filter cube (for the flow tracer), A4 filter cube (for nuclei), and DSR filter cube for the labeled protein.

The sections to be labeled with antibodies against ADAM15, chitinase 3-like 1 (CHI3L1), vimentin (VIM), estrogen receptor 1 (ESR1), and α-smooth muscle actin (α-SMA) were permeabilized with 0.2% Triton X-100 for 5 min and blocked in 2.5% donkey serum for 30 minutes at room temperature. Sections were then incubated with the primary antibody at room temperature for 3 hours, followed by incubation with Alexa Fluor 568 Donkey Anti-Rabbit secondary (1:1000) at room temperature for 60 minutes (see **Supplemental Table 1** for primary antibody details). Slides were counterstained with 2.5 µM Hoescht 33342 (Invitrogen) in TBS for 10 minutes to label nuclei, and glass coverslips were mounted with Vectashield Vibrance. Slides were imaged on a Nikon A1R confocal Ti2 microscope. Images were taken with a 20× objective plus an additional 2× intermediate magnification to obtain ND2 stack images with a 0.31 µm/pixel resolution. Nuclei were visible in the blue channel, the fluorescent tracer (Fluospheres) in the green channel, and the protein(s) of interest in the red or far-red channel. Images were then converted using ImageJ (FIJI) to maximum-intensity Z-projections and saved as BMP and TIFF files.

### 2.8. Immunofluorescence Quantification and Nuclei Counting

Quantification of fluorescent signal was performed using ImageJ (Fiji)^31^. The number of images quantified for each antibody target are specified in figure captions. In each section, a region of interest (ROI) was manually drawn around the TM using the polygon selection tool based on anatomical features visible in the brightfield channel (termination of Descemet’s membrane, TM beams and SC lumen) and the location of fluorescent tracer visible in the GFP channel. To quantify protein immunofluorescence, background correction was performed by subtracting the mean fluorescence measured in the sclera of each section, since we did not expect differences in scleral labeling between high vs. low flow regions for any of our target proteins. We then calculated the mean gray value of labeling for each antibody target within the TM ROI.

Due to the variability in tracer abundance in the designated HF and LF regions based on *en face* images, we also quantified tracer intensity within each tissue section, allowing us to use a continuous representation of segmental flow regions. In the tracer (GFP) channel, rolling-ball background subtraction (radius = 1.55 µm) was applied to isolate the tracer’s punctate fluorescent signal within the TM. The integrated density of tracer signal within the previously defined ROI was then quantified and used to represent the total tracer amount in the TM of each section.

Repeated-measures correlation (“rmcorr” package in R) was used to assess association between tracer and protein signals across tissue sections while accounting for non-independence of measurements from sections obtained from the same eye. A shared within-eye linear association between variables was estimated such that each correlation line shared the same slope while controlling for between-sample variability in baseline fluorescence (i.e., the y-intercept). For each protein marker, correlation coefficients (r_rm_) and p-values were reported.

In images labelled for myocilin, fibronectin, SPARC, collagen I, or collagen VI, the number of stained nuclei were also quantified using ImageJ (Fiji). The nuclei channel (DAPI) images were background corrected using a rolling ball radius of 30 pixels. Otsu thresholding was applied to isolate the relevant fluorescent signal, and nuclei within the TM ROI were counted using the “Analyze Particles” function, where particles over 50 pixels with a circularity between 0.25-1 were counted as nuclei.

## 3. Results

### 3.1. Differential Expression Gene Analysis in Human

In the human dataset, a total of 8,299 genes remained after removing low-expressed genes. We performed principal components analysis (PCA) before and after correcting for inter-donor variability (**Supplemental Figure 1**). Before correcting the data, all three donor eyes clustered separately, including the two eyes from the same donor, indicating that each eye exhibited a distinct transcriptional profile. Because the primary focus of this analysis is to compare high vs. low flow regions within the same eye, each eye (rather than each donor) was treated as an independent sample, i.e. each eye was treated as an independent biological replicate for downstream statistical analyses and when correcting for inter-eye variability with *limma,* which removed the eye-specific clustering. Because statistical analyses focus on within-eye differences rather than between donors, the inclusion of two eyes from the same donor does not confound the primary HF vs. LF comparison.

We used a linear model to compare transcriptomic profiles between ROIs collected from high and low flow regions in human eyes and identified 426 differentially expressed genes (p < 0.05). Of these genes, 249 were upregulated in HF regions, and 177 were upregulated in LF regions (**Figure 2A**). Hierarchical clustering of these 426 genes shows separation by flow region, and no clear clustering by donor within flow regions, suggesting that these differences were not specific to a single donor (**Supplemental Figure 2**).

**Figure 2:**
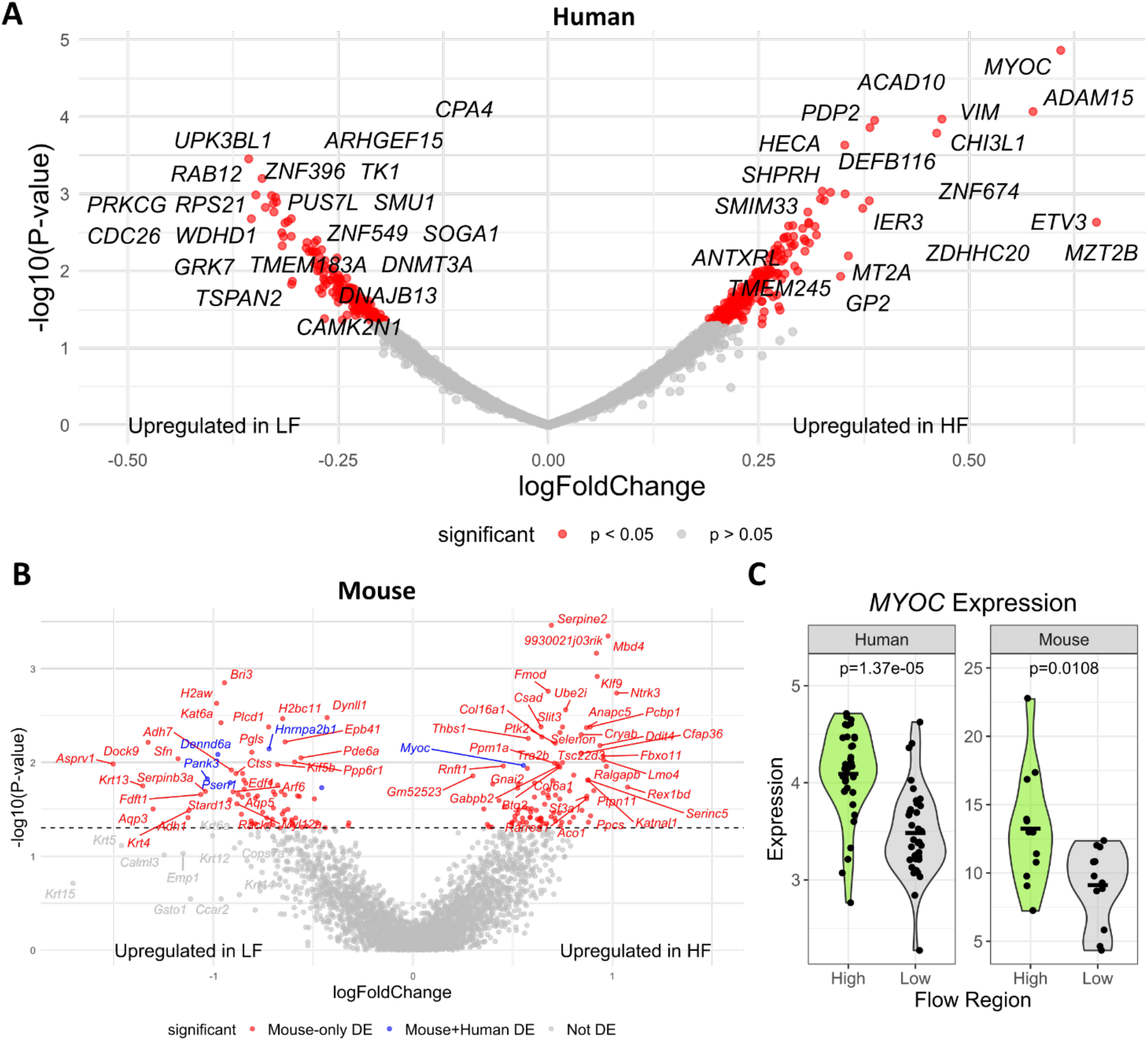
Differential Expression in Human and Mouse HF vs. LF TM. **(A)** Volcano plot showing differentially expressed genes in human TM. Each data point represents a single gene. Positive log fold change values indicate genes that had greater expression in HF regions, and negative values indicate greater expression in LF regions. Genes with p < 0.05 are shown in red. **(B)** Volcano plot showing differentially expressed genes in mouse TM. To highlight genes that overlap with the human differential expression analysis, genes with p < 0.05 <u>in mouse only</u> are shown in red, and genes with p < 0.05 <u>in both mouse and human</u> datasets are shown in blue. **(C)** Violin plots showing MYOC expression in both human and mouse HF vs. LF TM. In both species, MYOC was upregulated in HF regions (p < 0.05). Each point represents expression data from one ROI (Human: n=72 ROIs from 3 donor eyes; Mouse: n=24 ROIs from both eyes of 1 mouse).

One of the most significantly differentially expressed genes was myocilin (*MYOC*), a known glaucoma-related gene (logFC = 0.61, p = 1.37 × 10^-5^). Other top differentially expressed genes with known TM expression and/or glaucoma relevance that were upregulated in HF regions include ADAM metallopeptidase domain 15 (*ADAM15,* logFC = 0.58, p = 8.54 × 10^-5^), vimentin (*VIM, logFC = 0.47, p =* 1.07 × 10^-4^), and chitinase 3-like 1 (*CHI3L1, logFC = 0.46, p =* 1.63 × 10^-4^) (**Supplemental Figure 3**). A full list of the differentially expressed genes can be found in the Supplementary Files.

To verify that the transcriptomic data primarily represented TM cells, we assessed the expression of TM marker genes identified in a previous single-cell sequencing study of human conventional outflow pathway tissue^32^ (**Supplemental Figure 4**). We first examined canonical TM cell markers *MYOC*, *MGP*, *PDPN*, and *RARRES1*, all of which were detected above the limit of quantification (LOQ) in both HF and LF regions. We also assessed markers of previously identified Beam A, Beam B, and JCT TM cell subsets, including *FABP4* (Beam A), *TMEFF2* (Beam B), and JCT markers *CHI3L1*, *ANGPTL7*, *FMOD*, and *NELL2*. Although Beam A and B specific markers were not detected above the LOQ, three of the four JCT markers were detected in both HF and LF regions, further supporting the TM identity of the sequenced regions. We also did not detect key scleral markers, such as *PDGFRA* and *S100A4*, above the LOQ.

Because two of the most differentially expressed genes, *MYOC* and *CHI3L1*, are known markers of TM cells, we also performed nuclei counting in sections from HF and LF regions of the human donor eyes to confirm that expression-level changes were not due to differences in cellularity. We found no significant differences in nuclei count in HF vs. LF regions, indicating that these were in fact transcriptomic differences (**Supplemental Figure 5**).

We also examined the expression levels of known IOP- and POAG-associated genes in our dataset. Out of 228 genes containing significant IOP- and/or POAG-associated loci assembled from three genome-wide analyses^26–28^, 71 were detected above the LOQ in our dataset (**Supplemental Table 2**). Interestingly, six of those genes were significantly differentially expressed and upregulated in HF regions (**Table 2**).

**Table 2:** IOP- and POAG-associated genes differentially expressed between HF vs. LF regions. Six genes identified previously as significantly associated with IOP or POAG in genome-wide analyses were also differentially expressed between HF vs. LF regions. All six genes showed increased expression levels in HF regions.

| Gene | Genome-wide association | log Fold Change | P-Value |
| --- | --- | --- | --- |
| MYOC | IOP<sup>27</sup> & POAG<sup>26</sup> | 0.609018 | 1.37E-05 |
| ANTXR1 | IOP<sup>27,28</sup> & POAG<sup>26</sup> | 0.244433 | 0.015639 |
| ATXN2 | POAG<sup>26</sup> | 0.217422 | 0.02439 |
| BABAM2 | POAG<sup>26</sup> | 0.221136 | 0.026388 |
| OXR1 | IOP<sup>27</sup> | 0.211735 | 0.031857 |
| DAAM2 | IOP<sup>27</sup> | 0.234865 | 0.032465 |

### 3.2. Differential Expression Gene Analysis in Mice

We identified 167 differentially expressed genes in mouse HF versus LF TM regions (unadjusted p < 0.05), with 94 genes upregulated in HF regions and 73 upregulated in LF regions. Since we only have differential expression data from a single mouse, we focused on the genes that were also differentially expressed in our human data, since these are more likely to have biological significance (**Figure 2B**). Of the 167 mouse DE genes, six were also differentially expressed in the human dataset. Genes upregulated in LF regions in both species included *DENND6A*, *PANK3*, *HNRNPA2B1*, *PSEN,* and *SUB1*. Importantly, the only gene that was upregulated in both species in HF regions was *MYOC* (**Figure 2C**), further supporting a role for myocilin in segmental flow and outflow regulation.

### 3.3. Immunofluorescent Labeling of Top Differentially Expressed Genes in Human TM

We next performed immunofluorescent labeling of MYOC, CHI3L1, and ADAM15 proteins in human TM to investigate protein-level differences between HF vs. LF regions in our top differentially expressed genes. Contrary to our transcriptomic data, myocilin protein labeling was greater in LF regions than in HF regions (**Figure 3A, B**). Repeated measures correlation analysis in TM sections from two donor eyes (Donor 1 OD and Donor 2 OS) revealed a significant inverse correlation between myocilin labeling and tracer intensity (p < 0.01). This trend was consistent when labeled with primary antibodies sourced from two different commercial suppliers, including both a monoclonal and polyclonal antibody (**Supplemental Figure 6**), indicating that it is a robust finding. In addition to the TM proper, myocilin labeling was also present in the SC inner and outer wall in the LF sections. Conversely, CHI3L1 labelling was greater in the TM of HF sections compared to LF sections, agreeing with our transcriptomic data (**Figure 3C, D**). CHI3L1 immunofluorescent staining was significantly correlated with tracer intensity in the TM of both eyes from Donor 1 (p < 0.001). Note that, due to differences in imaging conditions (confocal vs. epifluorescence microscopy), absolute ranges in fluorescence intensity measurements differed between the correlation plots in Figures 3B and 3D. Even within plots, there is variability in fluorescence due to heterogeneity in tracer levels within regions classified as HF or LF, as well as inescapable variability in fluorescent protein labeling across different tissue sections. We believe that presenting plots of immunolabelling density vs. tracer fluorescence, as in Figure 3 B & D and subsequent figures, most transparently represents our data while still allowing correlations to be drawn between flow and protein levels.

**Figure 3:**
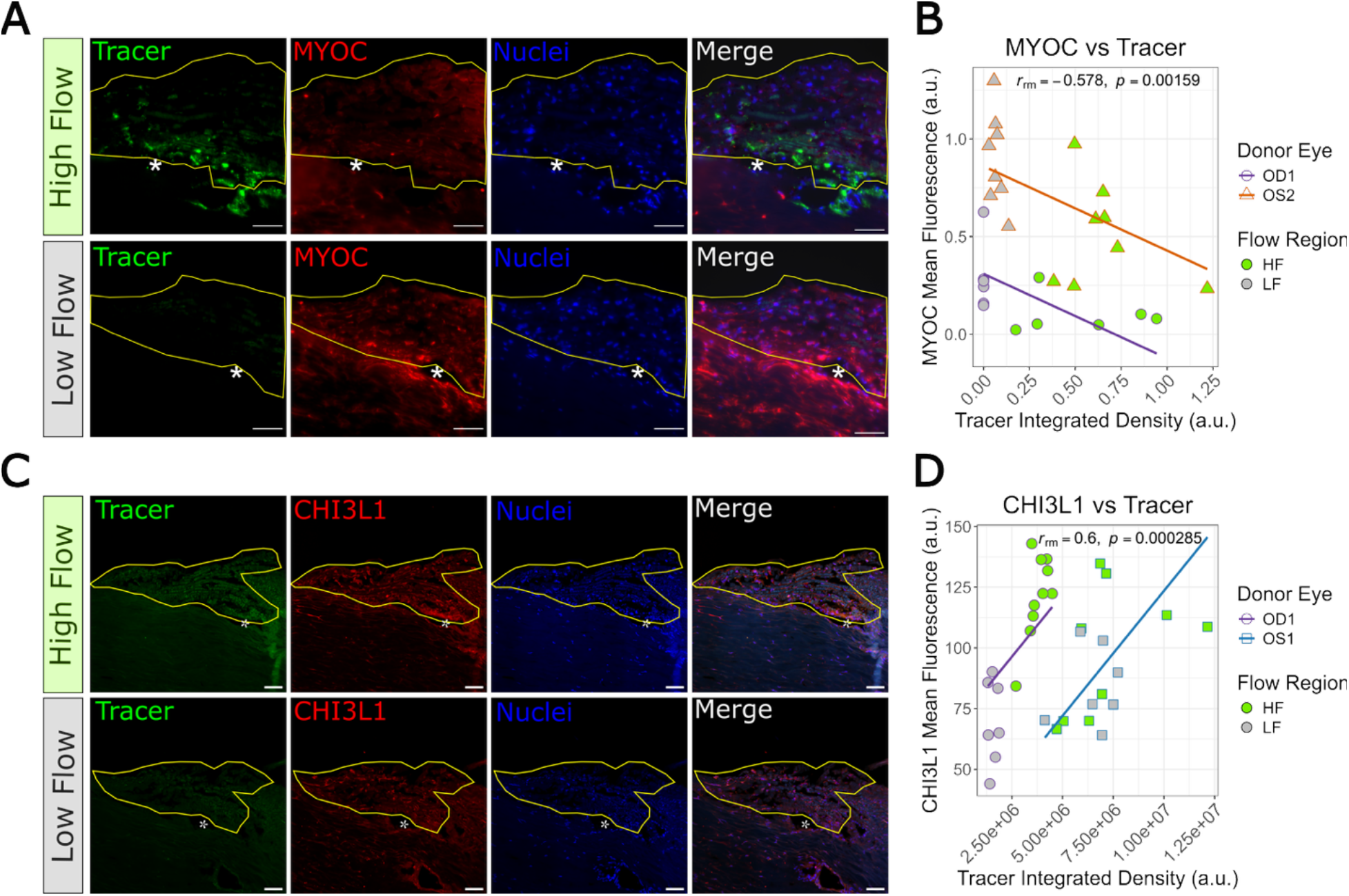
Immunofluorescent Staining of Myocilin and CHI3L1 in Human HF and LF TM. **(A)** Human TM sagittal sections (Donor 1, OD eye) from representative HF and LF regions labeled for myocilin. In all images, the TM is outlined in yellow, and the asterisk (*) denotes the SC lumen. Myocilin labeling was also visible in the inner and outer SC wall (surrounding the asterisk) in the LF sections. **(B)** Repeated-measures correlation plot showing association between MYOC labeling vs. tracer intensity in the TM in the OD eye of donor 1 (n=6 HF and 6 LF sections) and the OS eye of donor 2 (n=10 HF and 10 LF sections). Points represent individual sections, and parallel lines indicate donor eye-specific regression fits, with rrm being the correlation coefficient and p-values representing repeated measures correlation significance. **(C)** Human TM sagittal sections (Donor 1, OD eye) from representative HF and LF regions labeled with CHI3L1. **(D)** Repeated-measures correlation plots showing association between CHI3L1 labeling vs. tracer intensity in the TM in the OS and OD eye of donor 1 (n=10 HF and 8 LF sections per eye). Interpretation is described in Panel B. Note that axes scales differ between panels B and D due to differences in imaging acquisition conditions/microscope systems (see Methods). Scale bars = 50 μm.

We also labeled ADAM15 protein in TM sections and were able to detect it in both HF and LF regions; however, we did not observe significant correlation between ADAM15 and tracer intensity across all three donor eyes (**Supplemental Figure 7**). Together, these findings demonstrate that protein levels in different segmental flow regions only partially recapitulate the transcriptomic differences identified between HF vs. LF regions, highlighting the complexity of relating protein localization and transcript levels in the TM.

### 3.4. Human Gene Set Variation Analysis

We utilized gene set variation analysis (GSVA) to further analyze the transcriptomic count data and detect changes in groups of related transcripts in an unsupervised manner, allowing for investigation into pathway-level differences between HF and LF regions. Out of 5,482 total gene sets from the MSigDB2 C2 canonical gene set, 379 were found to be differentially enriched (p < 0.05) between HF and LF regions.

#### 3.4.1. Matrix-Related Differences in HF vs. LF Trabecular Meshwork

We first investigated differentially enriched gene sets related to ECM turnover, actin cytoskeleton, and integrins (**Figure 4**). Five gene sets were identified as being differentially enriched. Specifically, the “BIOCARTA ACTINY PATHWAY”, which contains genes related to Y-branching of actin filaments, and “REACTOME CELL EXTRACELLULAR MATRIX INTERACTIONS” gene sets showed greater enrichment in HF regions (p < 0.05), indicating upregulation of genes involved in actin filament remodeling and mechanosensory pathways in HF regions. Of the three gene sets that were enriched in LF regions, two were related to integrins, which also relate to cell-ECM interactions, and one was also related to actin remodeling. Together, this suggests that mechanosensation, cell-ECM interactions, and actin cytoskeletal remodeling differ between HF vs. LF regions of the TM.

**Figure 4:**
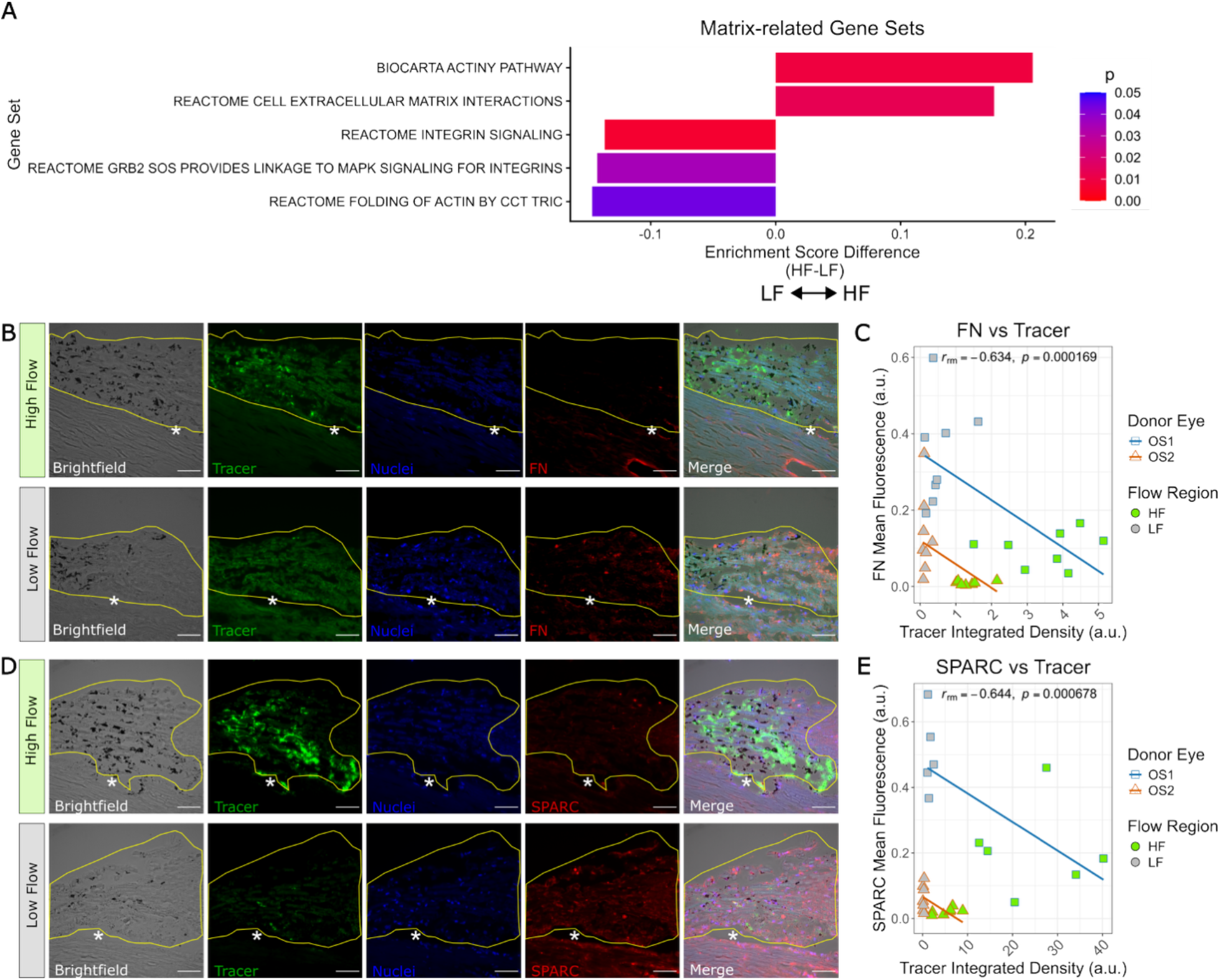
Matrix-related gene sets and protein levels differ between HF vs LF human TM. **(A)** Enrichment score differences (bars) for gene sets related to ECM, actin, and integrin signaling. Positive values indicate enrichment in HF regions, while negative values indicate enrichment in LF regions. Pathways labeled on the vertical axis are from the MSigDB C2 canonical gene set, and colors represent statistical significance. **(B)** Labeling of fibronectin in representative HF and LF human TM sections (Donor 2, OS eye). **(C)** Repeated-measures correlation plot showing association between fibronectin mean fluorescence intensity and tracer intensity in the TM for both Donor 1 OS eye and Donor 2 OS eye (n=8 HF and 8 LF sections from each eye). **(D)** SPARC labeling in representative HF and LF human TM sections (Donor 1, OS eye). **(E)** Repeated-measures correlation plot showing association between mean SPARC fluorescence intensity and tracer intensity in the Donor 1 OS eye (n=6 HF and 5 LF sections) and the Donor 2 OS eye (n= 7 HF and 7 LF sections). See Fig. 3 for interpretation of annotations on micrographs and correlation plots. Scale bars = 50 µm.

We also further investigated matrix-related differentially expressed genes to gain deeper insights into the transcripts driving the differential enrichment in actin remodeling and cell-ECM interactions (**Supplemental Figure 8**). Several top matrix related genes were expressed at higher levels in the HF regions, including microfibril associated protein 1 (*MFAP1*), TIMP metallopeptidase inhibitor 3 (*TIMP3*), insulin-like growth factor-binding protein 4 (*IGFPB4*), elastin (*ELN*) and versican (*VCAN*). One of the few ECM-related DE genes that was increased in LF regions was periostin (*POSTN*), a secreted ECM protein involved in cell adhesion.

Because we did not detect differences in many common TM ECM components at the transcript level, we decided to investigate whether there were matrix-related differences in HF and LF regions at the protein level, rather than the mRNA level, especially due to the differential expression of several matrix-modifying genes. We thus performed immunofluorescence microscopy labeling for fibronectin (FN), secreted protein acidic and cysteine rich (SPARC), and collagens I and VI, since these are important components of the TM ECM^5,33,34^. We observed greater FN and SPARC protein labeling in the LF regions compared to the HF regions in both donor 1 OS and donor 2 OS eyes (**Figure 4B-E**). Repeated-measures correlation analysis revealed a significant inverse correlation between fibronectin vs. tracer intensity (p < 0.001) and SPARC vs. tracer intensity (p < 0.001). While both collagens I and VI were present in the TM, there were no significant differences between flow regions (**Supplemental Figure 9**). Taken together, these results suggest that differences in fibronectin and SPARC levels between segmental flow regions are due to differences in ECM turnover and organization rather than gene expression.

#### 3.4.2. IOP-Relevant Pathways and Genes

In addition to the cytoskeletal and matrix-related gene sets, we also observed several differentially enriched gene sets likely related to IOP-associated genes. For example, three gene sets related to vascular endothelial growth factor (VEGF) were more enriched in HF vs. LF regions (**Figure 5A**). Specifically, these gene sets included genes that are targets of VEGF-A and are related to VEGF-A signaling. Many of the differentially expressed genes in these gene sets also appeared in the matrix-related gene sets, including *ELN*, *VCAN*, and *POSTN*. Notably, estrogen receptor alpha (*ESR1*) was a differentially expressed gene in the VEGF-related gene sets, with greater expression of *ESR1* in HF regions (**Figure 5D**). Despite increased VEGFA-related targets in HF regions, *VEGFB* was upregulated in LF regions (**Figure 5B**), and VEGFC was not differentially expressed between flow regions (**Supplemental Figure 10**). These results suggest potential differences in VEGF targets between high and low flow regions, as well as a potential role for estrogen signaling in segmental outflow.

**Figure 5:**
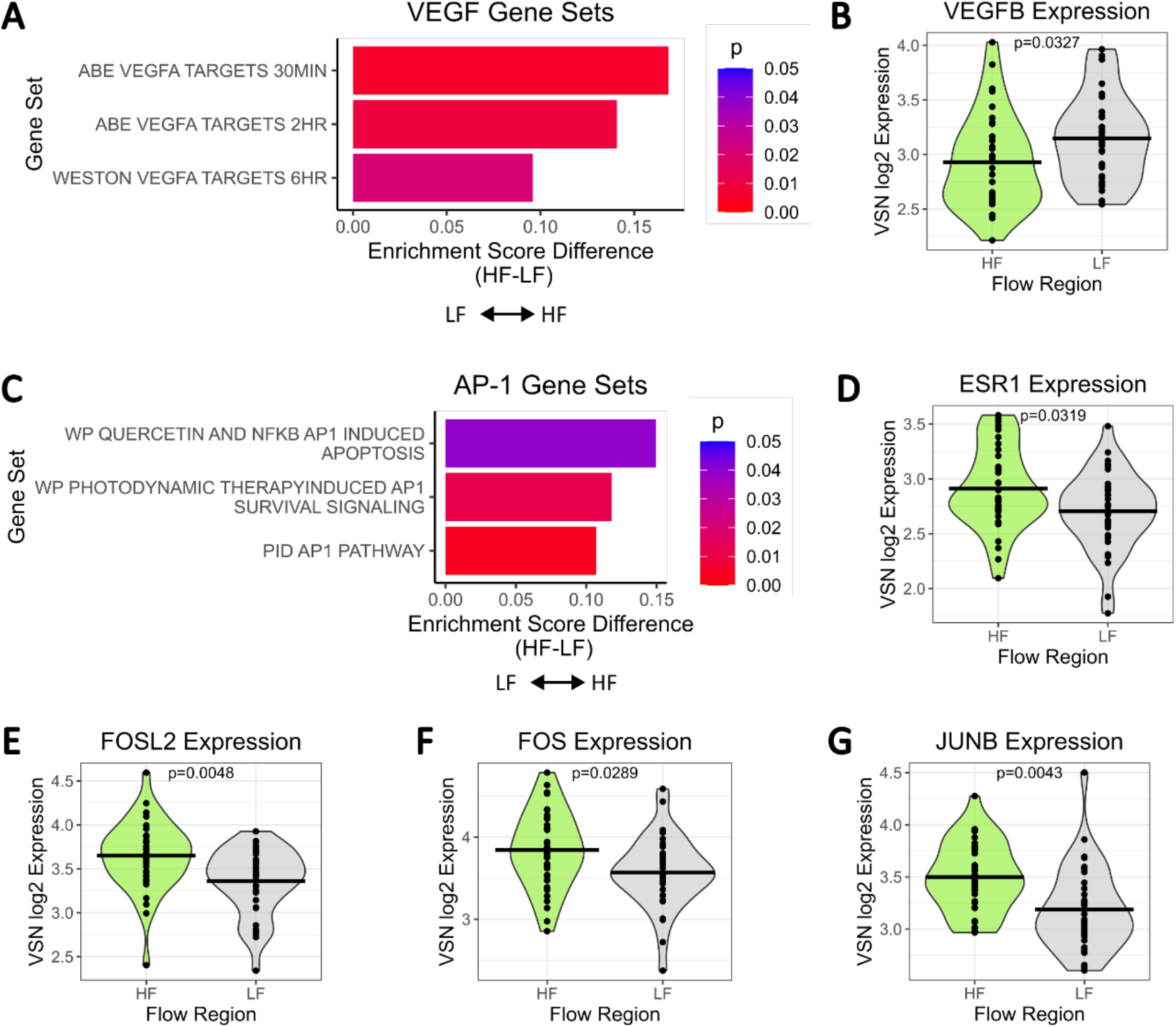
VEGF and transcription factor genes and gene sets. **(A)** Enrichment score differences for gene sets related to VEGF. Bars represent enrichment score differences. Positive values indicate greater expression of VEGFA targets in HF regions compared to LF regions. **(B)** Violin plot of normalized log2-transformed expression values of VEGFB, which was upregulated in LF regions. **(C)** Enrichment score differences for gene sets related to the AP-1 transcription factor. Positive values indicate greater expression of AP-1 components and related transcripts in HF regions compared to LF regions. **(D-G)** Violin plots of normalized (VSN) log2-transformed expression values of differentially expressed genes ESR1 (D), FOSL2 (E), FOS (F), and JUNB (G), which all showed greater expression in HF regions.

Gene sets associated with the activator protein-1 (AP-1) transcription factor were also among the most differentially enriched gene sets, with greater enrichment in HF regions relative to LF regions (**Figure 5C**). *ESR1* was also one differentially expressed gene included in these gene sets, along with *JUNB*, *FOS*, and *FOSL2* (**Figure 5E-G**). This pattern suggests that AP-1-mediated transcription may contribute to segmental outflow, potentially as a regulator of downstream gene expression differences between HF and LF regions. We performed immunofluorescent staining for ESR1 protein in sections from all three donor eyes, but no consistent correlation with tracer intensity was observed across donors (**Supplemental Figure 11**).

#### 3.4.3. Differences in Epigenetic Regulatory Gene Expression Between HF and LF Regions

We also noticed differential enrichment in gene sets related to epigenetic modifications, histone acetylation, and DNA methylation (**Figure 6**). Seven of these gene sets were enriched in HF regions, and four gene sets were enriched in LF regions. Specifically, several histone acetylation-related pathways, including “BIOCARTA HDAC PATHWAY”, “REACTOME HATS ACETYLATE HISTONES”, and “REACTOME HDACS DEACETYLATE HISTONES”, were enriched in HF regions. Notably, we also identified some complementary differences in gene sets between flow regions that related to specific histone deacetylases (HDACs). For example, “HDAC7 TARGETS 1 DN” was up in HF regions, while “HDAC7 TARGETS 1 UP” was up in LF regions, suggesting reciprocal regulation of HDAC7-associated targets between the two outflow regions. Similarly, HF regions showed greater enrichment of several methylation-associated gene sets (e.g., “REACTOME PRC2 METHYLATES HISTONES AND DNA”, “REACTOME PKMTS METHYLATE HISTONE LYSINES”, and “AML METHYLATION CLUSTER 4 UP”), while “MISSIAGLIA REGULATED BY METHYLATION DN” was enriched in LF regions, suggesting complementary differences in DNA methylation-related pathways between flow regions.

**Figure 6:**
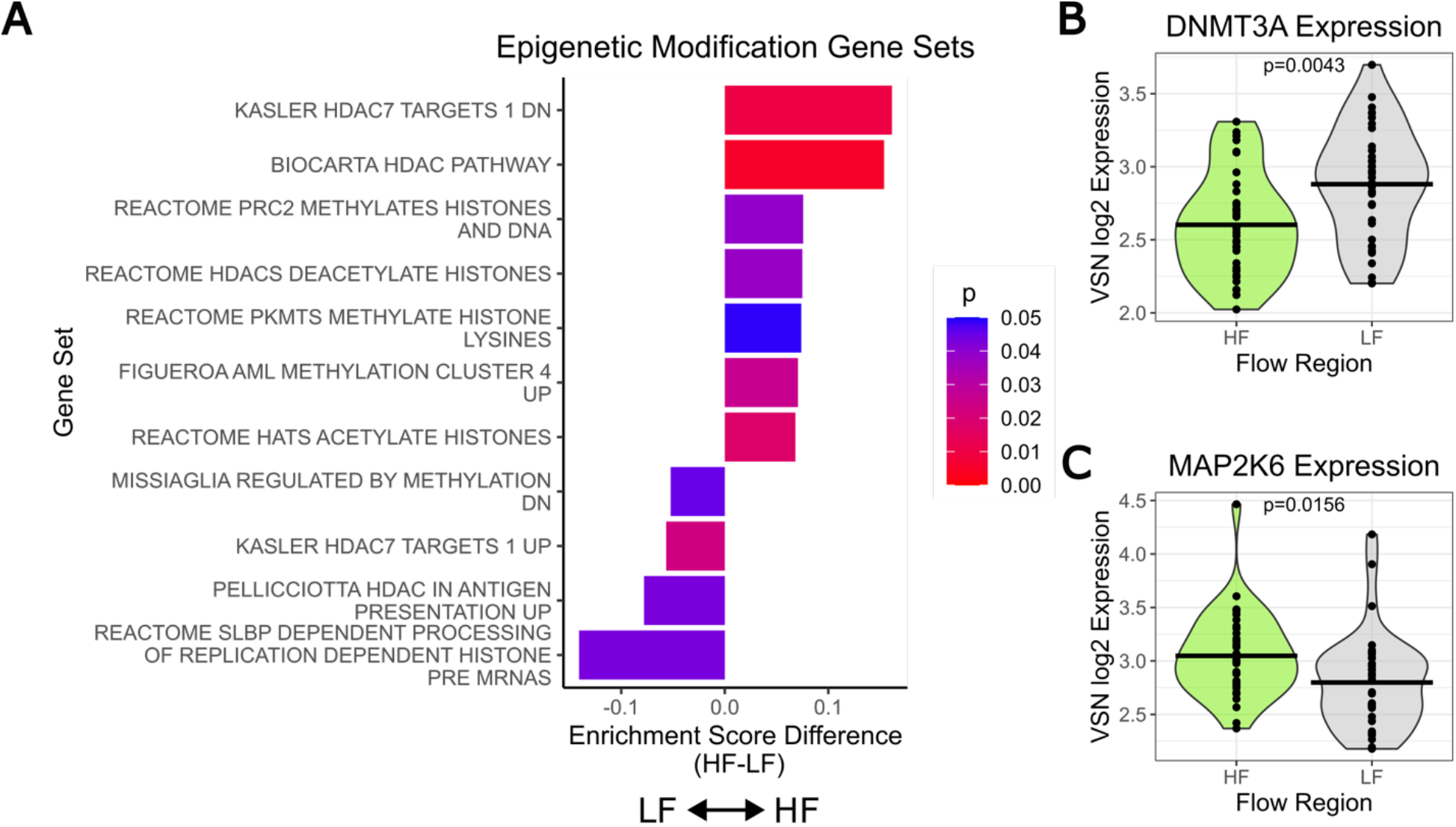
Epigenetic-Related Gene Sets are Differentially Enriched in HF and LF TM regions. **(A)** Eleven gene sets related to histone acetylation and DNA methylation were differentially enriched in HF vs. LF regions. Four gene sets were more enriched in LF regions, and 7 were enriched in HF regions, but pathways were complementary (i.e., “HDAC7 TARGETS1 UP” is up in LF, while “HDAC7 TARGETS1 DN” is up in HF). **(B)** Violin plot of normalized (VSN) log2 expression of DNMT3A, which is involved in DNA methylation and expressed more in LF regions vs. HF regions. **(C)** Violin plot of normalized log2 expression of MAP2K6, which is involved in histone acetylation and is expressed more in HF regions vs. LF regions.

Further examination of the differentially expressed genes in these gene sets supports complementary epigenetic modifications in HF vs. LF regions. For instance, *DNMT3A*, a DNA methyltransferase, was more highly expressed in LF regions (p < 0.01, **Figure 6B**), and *MAP2K6*, which is linked to histone acetylation, was more highly expressed in HF regions (p < 0.05, **Figure 6C**). Together, these results suggest a potential role for chromatin remodeling and epigenetic modification in segmental outflow. A full list of human GSVA gene sets with p < 0.05 is provided in the supplemental files.

### 3.5. Common Pathways Between Mouse and Human High and Low Flow Regions

To determine whether there were pathway-level transcriptional similarities between mouse and human TM, we next compared GSVA enrichment patterns between HF and LF regions across species. We identified three main groups of signaling pathways that were differentially enriched in both species’ HF vs. LF TM regions: Rho-kinase signaling, cellular stress pathways, and tumor necrosis factor-alpha (TNF-α) signaling.

#### 3.5.1. Rho-kinase Signaling is Enriched in LF Regions

GSVA revealed enrichment of RhoA- and Rho GTPase-related pathways in LF regions in both human and mouse TM (**Figure 7**). In the human data, only one RhoA-related gene set, “PID RHOA REG PATHWAY”, was significantly enriched in LF regions (p < 0.05). In mouse TM, six Rho GTPase-related gene sets showed greater enrichment in LF regions (**Figure 7A**). One Rho-associated gene set, “TRANFORMED BY RHOA DN”, was enriched in HF regions, but this gene set indicated decreased RhoA activity in HF regions relative to LF regions, which is consistent with the other gene sets. Overall, these findings suggest that RhoA/Rho-kinase (ROCK) signaling is elevated in LF regions of both mouse and human TM.

**Figure 7:**
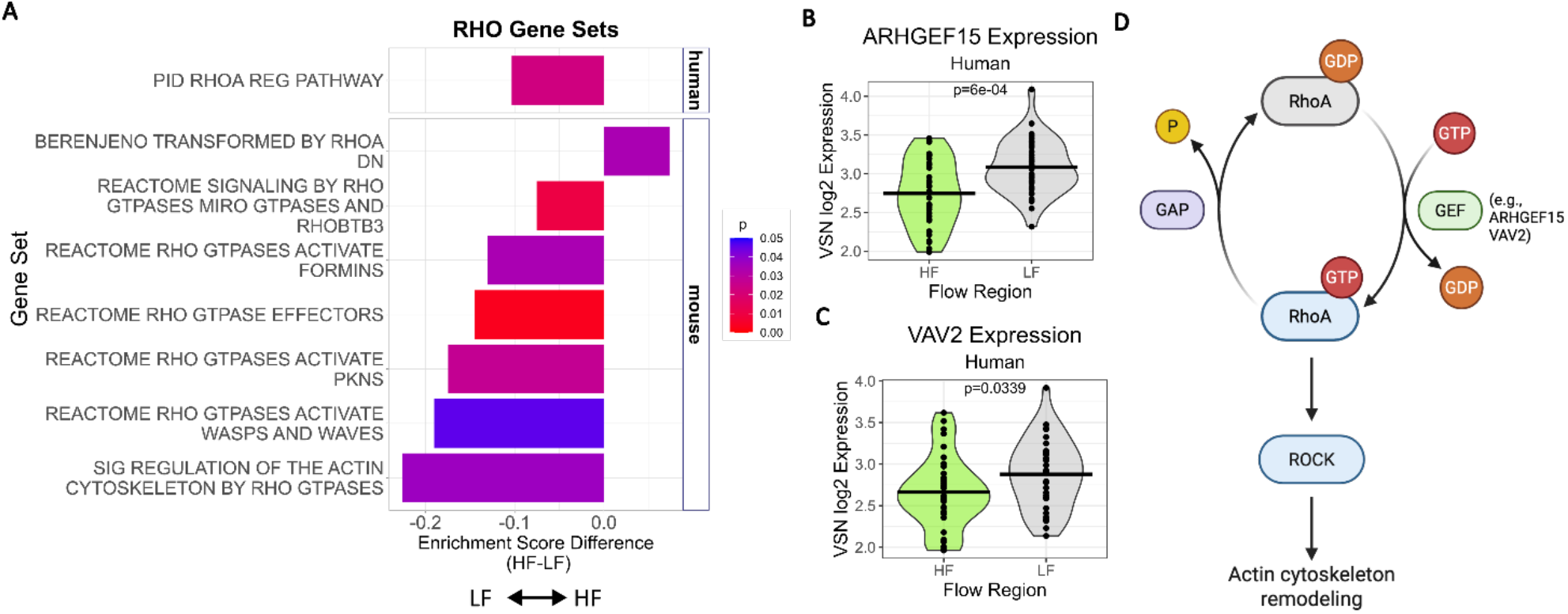
RhoA/ROCK Signaling is Enriched in LF Regions of Both Human and Mouse TM. **(A)** Enrichment score differences between HF vs. LF regions in RhoA and Rho-GTPase related gene sets in both the human and mouse datasets. In both species, gene sets related to RhoA signaling had greater enrichment in LF regions, except for “BERENJENO TRANSFORMED BY RHOA DN”, which was more enriched in HF regions. Because “RHOA DN” indicated downregulation of RhoA transformation, this suggests that overall, RhoA signaling is downregulated in HF regions. Only one gene set was differentially enriched (p < 0.05) in the human dataset, while 7 gene sets were significantly differentially enriched in the mouse dataset. **(B)** Violin plot of ARHGEF15 expression in human HF and LF regions. ARHGEF15, a Rho-A guanine exchange factor (GEF) that contributes to increased RhoA/ROCK pathway activation, was one of the most differentially expressed genes that was increased in LF regions in human TM. **(C)** Violin plot of VAV2 expression in human HF and LF regions. VAV2 is also a RhoA GEF that activates Rho/ROCK signaling. **(D)** Schematic of the RhoA/ROCK signaling pathway. Rho-A GEFs, such as ARHGEF15 and VAV2, promote RhoA/ROCK pathway activation which leads to actin cytoskeleton remodeling, increased cell stiffness, and decreased outflow facility in the TM^35^.

Although there was only one differentially enriched RhoA-related gene set in the human data, further inspection of individual genes within this set highlights its potential role in segmental flow. Notably, *ARHGEF15*, one of the most significant differentially expressed genes, was upregulated in LF regions and functions as a RhoA guanine nucleotide exchange factor (GEF), promoting RhoA–ROCK signaling (**Figure 7B, D**). Similarly, *VAV2*, another RhoA GEF upregulated in LF regions, further supports enhanced RhoA pathway activity in these regions (**Figure 7C, D**).

To interrogate differences in actin cytoskeleton remodeling at the protein level, we performed immunofluorescent staining of α-SMA in HF and LF TM sections from both human donors. Several LF sections appeared to have more α-SMA labeling compared to HF sections from the same eye, but the overall relationship between tracer and α-SMA staining did not reach statistical significance with repeated measures correlation analysis in sections from the OD and OS eyes from Donor 1 (**Supplemental Figure 12**).

#### 3.5.2. Cellular Stress Pathways and TNF-α Gene Sets are Enriched in HF Regions

We also identified similarities in enriched gene sets between humans and mice in pathways related to biological stressors such as hypoxia and reactive oxygen species (**Figure 8A**). In the human data, gene sets enriched in HF regions included one enriched gene set related to reactive oxygen species genes (“MOOTHA ROS”), two gene sets containing genes related to general oxidative stress response (“MIKHAYLOVA OXIDATIVE STRESS RESPONSE VIA VHL” and “WEIGEL OXIDATIVE STRESS RESPONSE”), and one containing genes related to oxidative stress-induced senescence (“REACTOME OXIDATIVE STRESS INDUCED SENESCENCE”). There were also two enriched gene sets related to hypoxia-inducible factor 1 subunit alpha (*HIF1A*), and four general hypoxia signaling gene sets. Notably, the *HIF1A* gene was not differentially expressed in the human data set (**Supplemental Figure 13**).

**Figure 8:**
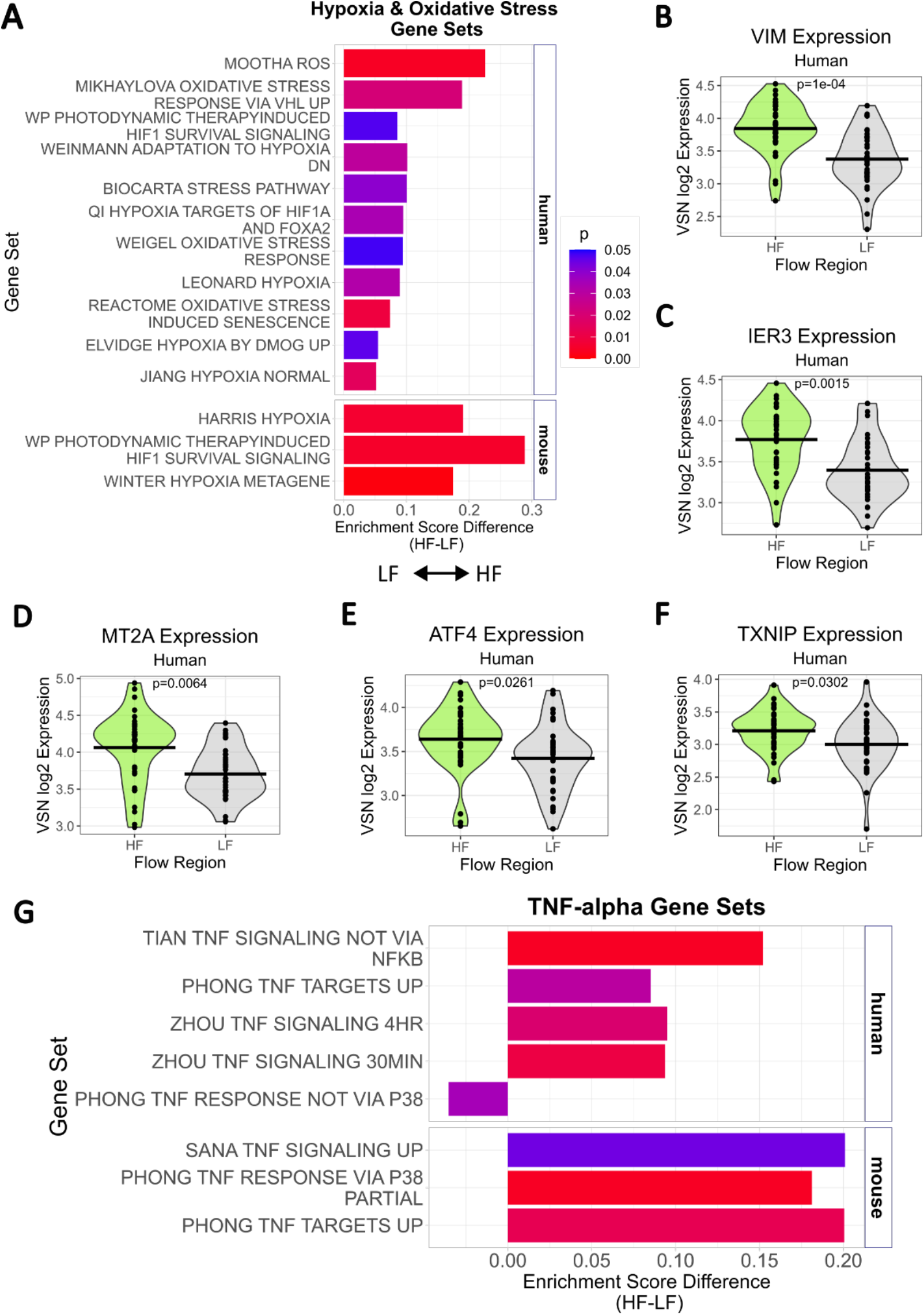
Cellular Stress-related and TNF-related Gene Sets are Enriched in HF Regions in both Human and Mouse TM. **(A)** Enrichment score differences in human and mouse gene sets related to oxidative stress, reactive oxygen species, hypoxia, and general stress response pathways. Eleven gene sets in human and three gene sets in mouse were more enriched in HF vs. LF regions. **(B-F)** Violin plots showing human expression of HF and LF differentially expressed genes from cell stress-related pathways. Vimentin (B), IER3 (C), MT2A (D), ATF4 (E), and TXNIP (F) were all expressed at higher levels in human HF TM compared to LF TM (p < 0.05). **(G)** Enrichment score differences in human and mouse gene sets related to TNF-α signaling. Of the five differentially enriched gene sets in the human TM, four showed greater enrichment in HF regions. Similarly, the three TNF-α signaling gene sets that were differentially enriched in the mouse TM were enriched in HF vs. LF regions. One specific gene set, “PHONG TNF TARGETS UP”, was significantly enriched in both human and mouse HF regions.

No oxidative-stress–related gene sets were differentially enriched in the mouse GSVA data; however, three hypoxia-related gene sets were significantly enriched in HF regions, consistent with the human data indicating increased expression of hypoxia-associated genes in HF TM tissue. We then investigated several of the specific genes differentially expressed in human HF vs. LF TM regions that were included in hypoxia and oxidative-stress related gene sets to better understand which genes may be driving some of the differences we observed at the pathway level (**Figure 8B-F**). Several stress-responsive genes were significantly upregulated in HF regions, including vimentin (*VIM),* an intermediate filament that remodels in response to cellular stress. We also performed immunofluorescent protein staining for vimentin in HF and LF human TM sections, but there was no significant correlation between tracer intensity and vimentin labeling across donor eyes (**Supplemental Figure 14**).

We also observed increased expression in HF vs. LF regions of Intermediate Early Response 3 (*IER3*), a mediator of resistance to stress and protection from TNF-α-induced apoptosis, and Metallothionein 2A (*MT2A)*, an antioxidant metallothionein that protects against oxidative damage. Additionally, activating transcription factor 4 (*ATF4)*, a key transcriptional regulator that mitigates endoplasmic reticulum (ER) stress, was upregulated, as was Thioredoxin-Interacting Protein (*TXNIP*), a modulator of oxidative and ER stress. Together, these findings indicate activation of multiple stress-response pathways in HF regions.

Of note, the BIOCARTA STRESS PATHWAY was enriched in HF regions in the human data (**Figure 8A**), and this pathway contains genes related to general and TNF-α stress related signaling. Thus, we further explored TNF-α signaling pathways in the human and mouse data sets. In concordance with our observations regarding cellular stress-related gene sets, both mouse and human GSVA data showed enrichment of TNF-α signaling pathways in HF regions (**Figure 8G**). Notably, the same “PHONG TNF TARGETS UP” gene set was enriched in HF regions in both human and mouse, suggesting a conserved TNF-α signaling pathway in segmental flow. Three other non-identical pathways that include genes related to TNF-α targets and signaling were also enriched in human HF regions, and one (“PHONG TNF RESPONSE NOT VIA P38”) was enriched in LF human TM, suggesting that the TNF-α signaling pathways that are upregulated in HF vs. LF regions may differ in their downstream targets. In mouse, two other gene sets related to TNF-α signaling and response were more enriched in HF regions, complementing the trends observed in human HF regions.

Interestingly, many of the differentially expressed TNF-α genes in the human data overlapped with stress-response genes (e.g., *IER3*) and transcription factors with known glaucoma/IOP-related roles (e.g., *JUNB*). Taken together, these findings point to a conserved TNF-α-associated stress response in HF TM cells that may influence local transcriptional regulation, ECM dynamics, and aqueous outflow dynamics (**Figure 9**). Overall, the increased expression of genes related to ROS and ER stress resistance, immune signaling, and transcription of *CHI3L1* and *MYOC*, which are known to be upregulated in response to various types of stress, suggest a more dynamic stress response in HF regions. Additionally, these regions show increased expression of ECM-related genes, *ADAM15*, and *VIM*, but decreased protein levels of FN and SPARC, indicating that there is increased ECM turnover in HF regions compared to LF regions, which have increased FN deposition at the protein level, but relatively lower expression of other ECM components at the mRNA level. Further, increased ROCK signaling in LF regions suggests that LF TM regions have more ECM accumulation and increased contractile tone. Human transcriptomic data suggests that histone modifications are distinct between HF and LF regions in humans, but further research is needed to confirm whether epigenetic modifications may allow HF regions to become LF, and vice versa.

**Figure 9:**
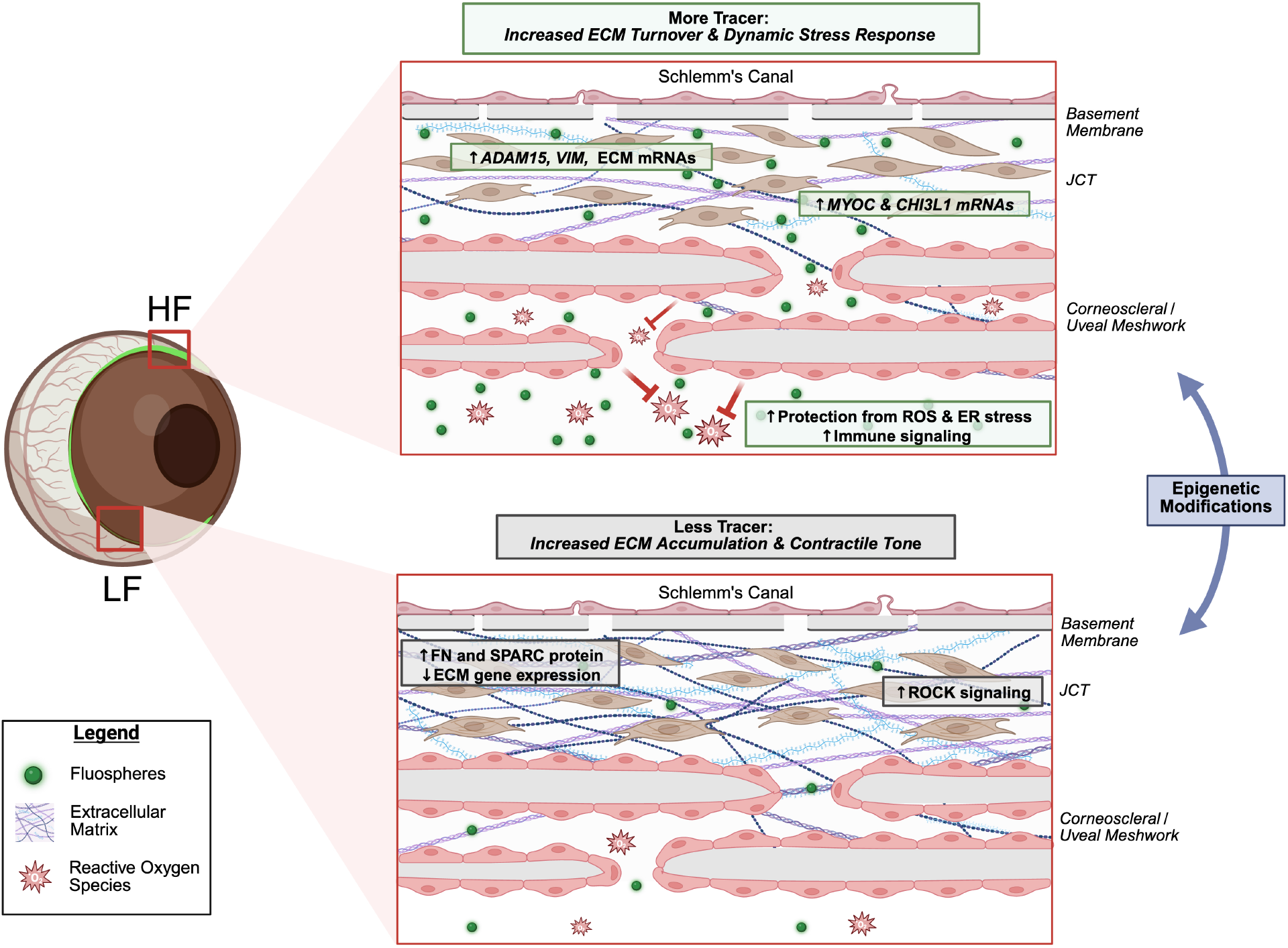
ECM Dynamics and Cell Stress Response in HF vs LF TM. High flow regions, indicated by a greater concentration of fluorescent tracer in the TM, have increased expression of ADAM15, VIM and other ECM-related mRNAs, and simultaneously show decreased protein labeling of ECM components compared to LF regions. HF TM cells also express MYOC and CHI3L1 in higher levels and have greater expression of genes related to protection from cellular stressors (i.e., oxidative, ER, immune stressors) compared to LF regions. Low flow regions, indicated by relatively low tracer in the TM, have lower ECM gene expression levels, but greater protein levels of ECM components, suggesting more ECM accumulation compared to HF regions. Cells in this region also have increased ROCK signaling, suggesting greater contractile tone in LF TM cells. Differences in histone acetylation and methylation pathways suggest that epigenetics may allow TM regions to transition between HF and LF gene expression patterns.

## 4. Discussion

Using spatial profiling of fixed frozen tissue sections from high and low flow regions of mouse and human TM, we identified and validated key differentially expressed genes and enriched gene sets, thereby enhancing our current understanding of segmental outflow.

While there have been previous studies investigating differentially expressed genes and proteins between HF and LF regions of human TM, the methods used here have identified differences in genes that have not previously been linked to outflow segmentation. Specifically, *MYOC* and *CHI3L1*, despite being well-studied in the context of TM biology, have not been previously identified as being differentially expressed in HF vs. LF regions. Most of the existing literature has focused on matrix-related genes and proteins using proteomic panels or selecting ECM-related transcripts for profiling with RT-PCR ^5,21,22^, which may not include genes such as *MYOC* and *CHI3L1*.

There has been one other study (Faralli et al.^20^) that used spatial profiling to interrogate the entire human transcriptome to compare HF vs. LF TM regions. Despite differences in the sets of differentially expressed genes identified between the two studies, the most significant previously reported differentially expressed gene (*ADAM15*) was also identified in our dataset. The differences in expression profiles reported in the two studies may be attributed to important methodological differences. Firstly, in the Faralli et al. study, the donor eyes were received up to 48 hours postmortem and perfused for several days in stationary organ culture before the Fluospheres perfusion and preparation for spatial profiling, whereas in our experimental setup, the perfusion of the beads was performed on whole globes within 5 hours postmortem. The difference in timing and perfusion methods may contribute to differences in detectable mRNA transcript levels. Secondly, the Faralli et al. study used paraffin-embedded tissue sections, while our study used fixed frozen sections, and this difference may also contribute to differences in which transcripts were preserved throughout tissue processing and/or could be detected by spatial profiling. Taken together, we hypothesize that these small but important differences in the experimental setup between the two studies contributed to the differences in transcripts detected, despite using the same transcriptomic profiling method.

### 4.1. Differentially Expressed Outflow- and Glaucoma-Related Genes

#### 4.1.1. Myocilin

One major finding of this study was the upregulation of myocilin in HF regions in both mouse and human TM. The relationship between *MYOC* mutations and ocular hypertension is well established, but the function of wild type myocilin is largely unknown. *MYOC* upregulation in response to dexamethasone treatment is a well-established phenomenon generally used as a marker for TM cells^36–38^. Several studies have reported *MYOC* upregulation in TM cells in response to stretch^39,40^; however, wild-type *MYOC* overexpression is not associated with increased IOP^41^, and *MYOC* expression is not required for normal IOP^42^. Higher levels of myocilin in the TM cause differential expression of genes related to cellular adhesion^43^, suggesting some interaction between myocilin and mechanical components of the tissue that influence outflow facility.

Somewhat surprisingly, myocilin protein labeling by immunofluorescence showed greater fluorescence intensity in LF regions of human TM tissue sections, despite higher transcript levels in the HF regions. In general, protein abundance can often show only modest concordance with transcript levels, with mRNA levels explaining as little as ∼20% of the variation in protein expression, highlighting the potential influence of post-translational regulation and in vivo protein stability^44–46^. Myocilin is a secreted protein, and differences in protein levels may be also due to differences in washout of secreted myocilin between HF and LF regions. Both antibodies we tested on our tissue target the N-terminal fragment, which is also “sticky” and may accumulate more in LF regions due to this property^47^, while it could more easily be washed out in regions of the TM with more outflow. This may partially explain why we observed more labeling by the antibody in LF regions, including in the SC inner and outer wall where *MYOC* expression is expected to be relatively low. Additionally, *MYOC* expression may be regulated by a feedback loop where low protein levels in the TM trigger upregulated transcription. The precise cause(s) of this apparent mismatch between transcript and protein levels in the TM is an intriguing area for future study.

#### 4.1.2. ADAM15

ADAM15, an integrin-binding metalloproteinase, was among the most significantly upregulated genes in HF regions, consistent with findings from Faralli et al.^20^. However, ADAM15 immunolabeling did not show a consistent protein-level trend across sections from all three donor eyes. Faralli et al. reported more labeling in HF regions in the JCT/SC inner wall region after separately quantifying ADAM15 in the TM beams, JCT/SC inner wall, and SC outer wall, suggesting that subregional differences in protein distribution might be present that were not captured by our whole-TM fluorescence quantification methods.

#### 4.1.3. CHI3L1

*CHI3L1* was also upregulated in HF regions of the human TM, and this finding was consistent at the protein level. *CHI3L1* is an established TM cell marker^48^ and is expressed at higher levels in normal TM cells compared to glaucomatous cells^49^, indicating a potential role in maintaining AH outflow homeostasis. *CHI3L1* transcription and secretion are also induced by TNF-α in TM cells^50^, which aligns with our GSVA results showing enriched TNF-α signaling in HF regions. *CHI3L1* is also upregulated in the TM under high-glucose conditions, indicating a potential role in response to cell stress^51^, further supporting the hypothesis that HF cells have an upregulated stress-response transcriptomic signature.

#### 4.1.4. Vimentin

We observed elevated expression of *VIM* in HF regions of the human TM, which aligns with its known role as a cytoskeletal component and regulator of cell stiffness in TM cells^52^. Vimentin has been reported to be elevated in glaucomatous and senescent TM cells ^49,52^, and it was included in oxidative stress pathways identified as enriched in HF regions in our GSVA results. While vimentin’s primary role as an intermediate filament is structural in nature, vimentin filaments are also known to rapidly remodel in response to oxidative stress^53,54^ as well as protein misfolding stress^55^. An increase in vimentin in HF regions may be in response to a greater metabolic load from more active filtering of AH outflow, as well as more mechanical stress compared to cells in LF regions. Additionally, vimentin has been shown to post-transcriptionally regulate synthesis and turnover of ECM components; collagen mRNAs associate with vimentin filaments, which prevents them from being translated^56–58^, which may explain why we did not observe differences in collagens between HF and LF regions. Previous studies have also reported that vimentin can help cells resist deformation in response to mechanical stretch^56^, suggesting that its upregulation may represent a response to higher outflow mechanical stress in HF regions. In summary, there are several potential mechanisms via which vimentin may contribute to maintaining TM cell integrity and thus outflow facility in a high flow environment.

#### 4.1.5. Transcription Factors

Several transcriptional regulators known to be associated with IOP homeostasis and AH outflow were upregulated in HF regions, including *ESR1*, *JUNB*, and *FOS*, which are all associated with the AP-1 transcription factor complex. This complex has also been shown to regulate *MYOC* expression^59,60^, aligning with the upregulation of *MYOC* in HF regions and positive enrichment of AP-1-related gene sets we observed in HF regions of human donor TM. Although AP-1 has been shown to upregulate *MMP3* in the TM^61^, we did not detect differential *MMP3* expression between HF and LF regions. We did, however, detect *TIMP3* upregulation in HF regions, which may reflect ESR1-associated modulation of the TIMP3/MMP axis^62^ and suggests active regulation of cell–ECM interactions and controlled matrix turnover in regions of higher outflow. Evidence also suggests that estrogen signaling via *ESR1* can be protective against elevated IOP and glaucoma risk^63^, and estrogen response signaling has also been shown to promote *MYOC* transcription in human TM cells^64^, consistent with our finding that both *ESR1* and *MYOC* were upregulated in HF regions. Taken together, the upregulation of transcription factors in HF regions suggests that segmental flow regulation, in part, may be regulated by modifying transcription of genes and proteins that influence outflow. We hypothesize that this may allow for homeostatic regulation of HF and LF TM regions and facilitate the dynamic nature of segmental flow.

#### 4.1.6. VEGF signaling

We detected increased enrichment of VEGFA-associated gene sets in HF regions of human TM, whereas *VEGFB* expression was higher in LF regions. VEGFA has been shown to increase outflow facility in mouse eyes and is stretch-activated in human TM cells^65^. Although *VEGFA* itself was not differentially expressed in our dataset, there was evidence that targets of VEGFA were expressed at higher levels in HF regions, suggesting that VEGFA signaling is more active in HF TM cells. Previous studies have demonstrated that VEGFA in the TM increases outflow facility, likely due to interactions with VEGF receptors in SC cells^65^. However, this relationship appears to be concentration dependent, as excessive VEGFA can trigger more fibrosis and actin crosslinking, both of which contribute to decreased facility^65,66^. Consistent with this, studies that did not differentiate between VEGF subtypes have reported that VEGF is increased in glaucomatous TM^67^. Notably, cells also upregulate VEGFA expression in response to mechanical and oxidative stress^65,68^, indicating that the observed differences in VEGF signaling may relate to the more dynamic stress-responsive transcriptional pattern observed in HF cells. Additionally, the upregulation of VEGFB in LF regions in our human data suggests that individual VEGF isoforms may play different roles in segmental outflow and, more broadly, in IOP homeostasis.

### 4.2. RhoA- and Rho-kinase-associated genes and gene sets

We observed increased RhoA/ROCK pathway enrichment in the LF regions of both human and mouse TM; further, in human tissue, we specifically identified two RhoA GEFs (*ARHGEF15* and *VAV2*) that were differentially expressed in the LF regions. This result is consistent with the established role of RhoA/ROCK signaling in modulating outflow resistance within the TM. Specifically, ROCK activation increases resistance to outflow by promoting actin stress fiber formation, cross-linking, and cytoskeletal reorganization, which in turn leads to synthesis and assembly of ECM components^69^. Thus, greater levels of RhoA GEFs in LF regions, which would increase ROCK activation, could lead to changes in cellular contractility and cytoskeletal/ECM phenotypes that are consistent with the LF regions having lower outflow facility and greater stiffness. Although ROCK inhibitors are already clinically used to lower IOP^70–72^ and are known to increase outflow through the TM^24^, their effect on segmental outflow remains poorly understood. Preferential targeting of LF regions may represent an opportunity for improved modulation of outflow facility, and this could be a potential focus for future studies.

### 4.3. Role of Cellular Stress in High and Low Flow Regions

We observed enrichment of hypoxia, oxidative stress, and TNF-α-related gene sets in HF regions. All these pathways could be considered as stressors to the TM cells, and increased enrichment of genes related to cell stressors is consistent with HF areas experiencing more metabolic stress due to higher AH outflow and increased exposure to materials transported in the aqueous humor, such as ROS, pigment, and other signaling molecules or cytokines. Consistent with the pathway-level findings, three of the differentially expressed genes identified from POAG GWAS studies, *OXR1*, *BABAM2*, and *ATXN2*, are associated with cellular stress responses and showed increased expression in HF regions. Specifically, *OXR1* is associated with a resisting oxidative stress^73,74^, *BABAM2* is associated with TNF-alpha signaling and protection from DNA damage in response to cellular stress^75,76^, and ATXN2 is involved in stress granule formation and general protection from cellular stressors^77,78^. Together, this further implicates the role of stress-response pathways in the TM in maintaining outflow homeostasis.

Interestingly, increased TNF-α signaling in HF regions is consistent with our current understanding of the mechanisms of selective laser trabeculoplasty (SLT), a clinical procedure that can enhance outflow facility and lower IOP in glaucoma patients. SLT induces inflammatory cytokine release, including TNF-α, either directly from TM cells or via monocyte/macrophage recruitment, which triggers ECM remodeling within the TM and potentially the SC inner wall to restore outflow homeostasis^79–82^. In the context of segmental outflow, greater expression of TNF-α downstream signaling targets by TM cells in HF regions may reflect greater exposure to inflammatory cytokines or other immunomodulatory signals carried by the aqueous humor, although factors such as local flow velocities and ligand-receptor kinetics may make the actual degree of exposure difficult to infer directly.

While hypoxia-related gene sets were enriched in HF regions, it is not clear whether these regions of the TM are truly more hypoxic. Notably, we did not observe any differential expression of the hypoxia marker *HIF1A* at the transcript level, and thus genes included in significantly enriched gene sets may instead reflect activation of related downstream stress-responsive signaling pathways. For example, *MAP2K6* is one gene contributing to hypoxia and cell stress-related pathways. Although it was first mentioned in our results as a histone acetylation-related gene, *MAP2K6* is also activated by hypoxic stress^83^, and it can also be induced by inflammatory stressors and TNF-α or environmental stress via integrin-mediated signaling. *MAP2K6* is part of the MAPK signaling cascade, which intersects with transcription factors such as *JUNB* and *FOS*, which were also differentially expressed in our data. These signaling pathways are also linked to downstream regulation of cytoskeletal components, cell contractility, and ECM remodeling, and stress-responsive processes more broadly. Together, these observations reinforce the idea that multiple stress-responsive pathways may converge to coordinate transcriptional, structural, and cell-ECM interactions that enable HF TM cells to maintain their function in a high outflow environment.

### 4.4. Epigenetic modification as a potential mediator of segmental outflow

Several gene sets related to chromatin remodeling, DNA methylation, histone acetylation and deacetylation differed between HF and LF regions, suggesting that epigenetic modification may contribute to the establishment and/or maintenance of segmental behavior in TM cells from HF and LF regions. This idea is consistent with prior observations that HF- and LF-derived TM cells retain distinct phenotypes in cell culture, even after being removed from the HF or LF environment in the eye^16,21,23^. Relatedly, it has been shown that segmental outflow is dynamic in young mice and regions can change from high to low flow over the course of 14 days; however, in older mice, the regions are less dynamic^10^. When HF and LF cells are extracted from human donor eyes, typically they are from older individuals. These observations, combined with the transcriptomic results from this study, suggest that epigenetic programming, potentially in conjunction with real-time exposure to AH outflow, may contribute to HF or LF phenotypes in TM cells, and these phenotypes become more stable with aging.

### 4.5. Limitations

In this study, we primarily focused on genes and proteins that were differentially expressed in human samples, or that were similarly differentially expressed across both human and mouse samples. The mouse data were interpreted more cautiously, as sequencing was only done from two eyes of a single mouse due to the challenging experimental setup and technical constraints with the GeoMx Spatial Profiling system, in addition to the small number of cells captured in each ROI even after combining the TM from two adjacent sections, it is likely that our ability to detect lower-abundance transcripts was limited. As a result, our mouse analysis may be biased towards genes with relatively high baseline expression levels.

Even in the human dataset, statistical power was limited due to the small sample size. It is therefore not surprising that no genes met an FDR-adjusted threshold of less than 0.05, given that only three donor eyes were available and successfully perfused to label flow regions. Nonetheless, we considered the unadjusted p-values valuable as exploratory indicators of potential differences between flow regions that can guide more targeted validation studies with larger cohorts in future studies. Additionally, we anticipated relatively subtle biological differences between HF and LF regions from an otherwise healthy TM, and the small log-fold change values of the differentially expressed genes supports this hypothesis. Because these donors did not have glaucoma or any other known ocular pathologies, the TM was likely able to maintain AH homeostasis, and any transcriptional differences between HF and LF regions were expected to have a modest effect size after adjusting for multiple comparisons. However, these subtle differences in gene expression between HF and LF regions are evidence that small changes in the expression profile of TM cells can potentially have a dramatic impact on local outflow resistance. This has important implications for our understanding of POAG disease mechanisms, as well as for the development of IOP-lowering therapeutics targeting TM cells.

The age difference between human and mouse eyes used here is also an important consideration. The human donor eyes were relatively old (72 and 73 years), while the mouse eyes were obtained from an 8-month-old animal, which corresponds to middle-age in mice^84^. Certain regulatory mechanisms for segmental outflow may become more stable and/or prominent with age, and it is possible that some transcriptomic differences, in epigenetic regulation for instance, may have only been detectable in the human dataset and not fully established in the relatively younger mouse eyes. Further, the fact that human donor samples were necessarily post-mortem tissue may have triggered elevated expression levels of hypoxia-related genes due to lower oxygen levels. It is possible that the HF and LF cells responded differently to the low-oxygen environment, and this is an important consideration for future studies using post-mortem donor tissue.

Finally, it is important to note that transcriptional differences identified here cannot determine causality; in other words, it cannot be determined from this dataset whether differences in specific transcripts drive HF or LF behavior, or if they arise as responses to local differences in aqueous humor outflow rate. Additionally, the function and transcription signatures of the corneoscleral and uveal meshwork, the JCT, and the SC inner wall are all slightly different, and we did not separate contributions from these subpopulations of cells that were encapsulated in our regions of interest for sequencing. Future studies using other approaches will be needed to determine whether the observed transcriptomic differences are upstream determinants or downstream consequences of segmental outflow.

## 5. Conclusions

By applying GeoMx spatial profiling to fixed frozen tissue sections taken from high and low flow regions of human and mouse trabecular meshwork, we generated the first cross-species transcriptomic study of segmental outflow. The identification of overlapping features, such as myocilin expression and broader pathway-level similarities across species, provides novel insights into the underlying biology of segmental outflow.

Our data further implicated genes linked to segmental outflow (e.g., *ADAM15*) and aligns with existing knowledge of genes and signaling pathways that are relevant in outflow dynamics and IOP homeostasis, including cell-ECM interactions and VEGF signaling. HF regions showed transcriptional signatures associated with inflammatory, environmental, hypoxic, and oxidative stress-responses, as well as increased expression of major transcription factors (e.g., *ESR1*, *JUNB*, *FOS*). LF regions showed higher expression of rho-kinase signaling and cytoskeletal remodeling-related genes and more ECM accumulation. We also identified differential enrichment of genes and gene sets related to chromatin remodeling, suggesting a potential contribution of epigenetic modification to segmental outflow (**Figure 9**).

Importantly, our protein-level validation with immunofluorescence microscopy highlighted that transcript levels and protein abundance are not always concordant; for example, fibronectin and SPARC showed increased protein levels in LF regions, but no difference at the transcriptomic level, while some of our top differentially expressed genes did not exhibit any differences between flow regions at the protein level. These findings underscore the necessity of integrating transcriptomic, proteomic, and functional differences in segmental flow regions in future studies.

## Supporting information

Human Differential Expression Data

Human GSVA data

Mouse Differential Expression data

Mouse GSVA data

## Acknowledgements

The authors thank the Technology Access Program at Bruker Spatial Biology for their assistance with this study. The authors also acknowledge the following funding sources: NIH T32 GM145735 [CAW], NIH EY031710 [CAW, CRE, WDS], Georgia Research Alliance [CRE], NIH R01EY028608, R01 EY022359, Research to Prevent Blindness Departmental Grant, and P30EY005722 [WDS], Challenge Grant from Research to Prevent Blindness, Inc. to the Department of Ophthalmology at Emory University, and NIH P30EY06360 (to the Atlanta Vision Research Community).

## Commercial Relationships Disclosure

C.A. Wong, None; A. T. Read, None; M. Chrenek, None; G. Li, None; W.D. Stamer, None; L.B. Wood, None; C.R. Ethier, None

