## Supplemental Data for "Transcriptomic Profiling of High vs. Low Flow Regions of Mouse and Human Trabecular Meshwork"

### 1. Supplementary Data

*Methods: Mouse Tissue preparation and GeoMx Spatial Profiling*

All mouse tissue processing was performed using RNAse-free protocols. Whole globes were cryoprotected, embedded in optimal cutting temperature compound (OCT), snap frozen in 2-methylbutane cooled with liquid nitrogen, and stored at −80 °C. Cryosections from high and low flow regions (8 μm thick) were collected using a CryoStar NX70 cryostat (Thermofisher) as described for the human tissue, and sections were arranged on Superfrost Plus Gold slides (Fisher) to facilitate processing in the GeoMx platform.

Two slides were carefully curated for processing in the GeoMx platform. On each slide, we included sections from both high and low flow, as well as OD and OS eyes. Based on the GeoMx platform guidelines recommending inclusion of approximately 50–100 cells per ROI (with optimal sensitivity achieved near 100 cells), tissue sections were arranged on the slide in pairs (**Figure 1D**). Each pair consisted of two adjacent sections positioned such that a single ROI could encompass the target tissue (TM) from both sections. This arrangement was designed to maximize the number of cells captured within each ROI, thereby improving our ability to detect more transcripts.

Four pairs of sections from each experimental group (i.e., high and low flow from each of the OD and OS eyes) were included on a single slide. Sections were also trimmed under a dissecting microscope so that the ROI could be drawn without intersecting the surrounding cornea, iris, ciliary body, or displaced retinal tissue (**Figure 1E**). Then, sections were allowed to dry and stored at −80 °C. Slides were shipped on dry ice to the Technology Access Program (TAP) team and processed according to standard workflows as described above. The slides were hybridized overnight with the GeoMx Mouse Whole Transcriptome Atlas (WTA) nucleic acid probes coupled with photocleavable oligonucleotide tags, and they were also labeled with primary antibody morphology markers against $\alpha$-SMA (Abcam, ab184675, 2.50 $\mu$g/ml) and CD31 (R&D Systems, AF3628, 3.33 $\mu$g/ml), and the 3 pairs of sections with the most intact morphology after slide preparation were chosen for ROI selection so that 12 ROIs total could be sequenced on each slide, resulting in a total of 24 ROIs in our dataset.

***Supplemental Table 1: Primary Antibodies used for Immunofluorescent Labeling.*** *Myocilin labeling using the Abcam Systems primary antibody is reported in the main figures, and the R&D Systems primary antibody labeling can be found in **Supplemental Figure 6**.*

| Target | Source | Catalog No. | Dilution |
|---|---|---|---|
| **Fibronectin** | Abcam | ab2413 | 1:500 |
| **SPARC** | Proteintech | 15274-1-AP | 1:500 |

| | | | |
|---|---|---|---|
| **Collagen I** | Invitrogen | PA1-26204 | 1:100 |
| **Collagen VI** | Novus Biologicals | NB120-6588 | 1:200 |
| **Myocilin** | R&D Systems | MAB3446 | 1:200 |
| | Abcam | ab41552 | 1:200 |
| **CHI3L1** | Invitrogen | PA5-43746 | 1:100 |
| **ADAM15** | Abcam | ab124698 | 1:500 |
| **VIM** | Santa Cruz | sc-7557 | 1:1000 |
| **ESR1** | Santa Cruz | sc-787 | 1:50 |
| **alpha-SMA** | Sigma | C6198 | 1:400 |

***Supplemental Table 2: Differential expression of IOP- and POAG-associated genes detected in HF and LF regions.*** *Genes with p<0.05 are highlighted.*

| Gene | Detected | logFC | P.Value | adj.P.Val |
|---|---|---|---|---|
| MYOC | TRUE | 0.609018 | 1.37E-05 | 0.113626 |
| ANTXR1 | TRUE | 0.244433 | 0.015639 | 0.84027 |
| ATXN2 | TRUE | 0.217422 | 0.02439 | 0.874591 |
| BABAM2 | TRUE | 0.221136 | 0.026388 | 0.88089 |
| OXR1 | TRUE | 0.211735 | 0.031857 | 0.887596 |
| DAAM2 | TRUE | 0.234865 | 0.032465 | 0.889196 |
| RAPSN | TRUE | 0.181084 | 0.072347 | 0.943735 |
| CTTNBP2 | TRUE | 0.159059 | 0.106319 | 0.943735 |
| DLL1 | TRUE | 0.226161 | 0.118642 | 0.943735 |
| VEGFC | TRUE | -0.15272 | 0.127262 | 0.943735 |
| SPTBN1 | TRUE | 0.154917 | 0.138622 | 0.945545 |
| ARHGEF12 | TRUE | -0.14608 | 0.154192 | 0.945545 |
| TFEC | TRUE | 0.142253 | 0.160658 | 0.945545 |
| BCAS3 | TRUE | 0.144761 | 0.161015 | 0.945545 |
| HGF | TRUE | -0.14683 | 0.162662 | 0.945545 |
| GLIS1 | TRUE | -0.13476 | 0.171722 | 0.945545 |
| SPTSSA | TRUE | 0.132417 | 0.177969 | 0.949007 |
| GAB2 | TRUE | -0.12139 | 0.230132 | 0.962433 |
| RPLP2 | TRUE | 0.127758 | 0.24758 | 0.962433 |
| ARHGEF3 | TRUE | -0.11734 | 0.249818 | 0.962433 |
| LOXL1 | TRUE | -0.10939 | 0.262877 | 0.962433 |
| ANGPT2 | TRUE | 0.113931 | 0.269423 | 0.962433 |
| CADM1 | TRUE | 0.1029 | 0.306182 | 0.962433 |
| ANKH | TRUE | 0.098817 | 0.312543 | 0.962433 |
| SCFD2 | TRUE | 0.103649 | 0.314832 | 0.962433 |

| | | | | |
|---|---|---|---|---|
| STOX2 | TRUE | 0.096991 | 0.317076 | 0.962433 |
| ZNF652 | TRUE | -0.09884 | 0.32362 | 0.962433 |
| SOS2 | TRUE | -0.09811 | 0.33171 | 0.962433 |
| TCF7L2 | TRUE | 0.085955 | 0.387947 | 0.967433 |
| YAP1 | TRUE | 0.092229 | 0.394528 | 0.967433 |
| CAPZA1 | TRUE | -0.08142 | 0.418903 | 0.969928 |
| FBXO32 | TRUE | 0.084957 | 0.424778 | 0.969928 |
| SRSF3 | TRUE | -0.08188 | 0.431855 | 0.969928 |
| SCAMP1 | TRUE | -0.08105 | 0.433345 | 0.969928 |
| UBIAD1 | TRUE | -0.07782 | 0.435114 | 0.969928 |
| RALGPS1 | TRUE | 0.0756 | 0.460069 | 0.972378 |
| ADAMTS18 | TRUE | -0.06894 | 0.476901 | 0.972378 |
| TMCO1 | TRUE | 0.070309 | 0.483358 | 0.972378 |
| CDH11 | TRUE | 0.073207 | 0.49073 | 0.973789 |
| SVEP1 | TRUE | 0.061234 | 0.528234 | 0.980499 |
| EFEMP1 | TRUE | 0.062429 | 0.531733 | 0.980639 |
| MYOF | TRUE | 0.061417 | 0.542278 | 0.980802 |
| RSPO1 | TRUE | 0.057447 | 0.546486 | 0.980817 |
| FNDC3B | TRUE | 0.063842 | 0.548569 | 0.980817 |
| AFAP1 | TRUE | 0.064702 | 0.55348 | 0.981257 |
| PTPRJ | TRUE | -0.06017 | 0.554368 | 0.982182 |
| PLCE1 | TRUE | -0.0606 | 0.566683 | 0.982182 |
| SYN3 | TRUE | 0.05351 | 0.605021 | 0.982182 |
| SYT13 | TRUE | -0.04343 | 0.670426 | 0.986027 |
| GTF2E2 | TRUE | -0.04231 | 0.676172 | 0.98654 |
| NPEPPS | TRUE | 0.039114 | 0.687283 | 0.987305 |
| CREB5 | TRUE | 0.041814 | 0.690776 | 0.987305 |
| CADM2 | TRUE | -0.03912 | 0.695219 | 0.987305 |
| TNS1 | TRUE | -0.03962 | 0.698969 | 0.987305 |
| PLEKHA7 | TRUE | -0.03468 | 0.728907 | 0.991294 |
| COL11A1 | TRUE | 0.034946 | 0.730721 | 0.991294 |
| RERE | TRUE | 0.033192 | 0.731748 | 0.991294 |
| ANAPC1 | TRUE | 0.032832 | 0.753348 | 0.993209 |
| HHEX | TRUE | 0.030198 | 0.765005 | 0.993209 |
| RARB | TRUE | -0.02567 | 0.797245 | 0.993209 |
| FOXC1 | TRUE | 0.026538 | 0.797743 | 0.993209 |
| TMEM181 | TRUE | 0.024107 | 0.807747 | 0.993209 |
| THSD7A | TRUE | 0.020352 | 0.843808 | 0.993209 |
| FERMT2 | TRUE | -0.01785 | 0.856593 | 0.993209 |
| LHPP | TRUE | 0.015349 | 0.877705 | 0.993209 |
| TES | TRUE | 0.008515 | 0.93032 | 0.994798 |

| | | | | |
|---|---|---|---|---|
| CLIC5 | TRUE | 0.008429 | 0.930509 | 0.994798 |
| TLL1 | TRUE | 0.008938 | 0.931418 | 0.994798 |
| IGF1 | TRUE | 0.008405 | 0.933485 | 0.994798 |
| MAPT | TRUE | -0.00885 | 0.9354 | 0.994798 |
| NR1H3 | TRUE | -0.00485 | 0.961184 | 0.996064 |
