## Supplemental Figures for "Transcriptomic Profiling of High vs. Low Flow Regions of Mouse and Human Trabecular Meshwork"

### 1. Supplemental Figures

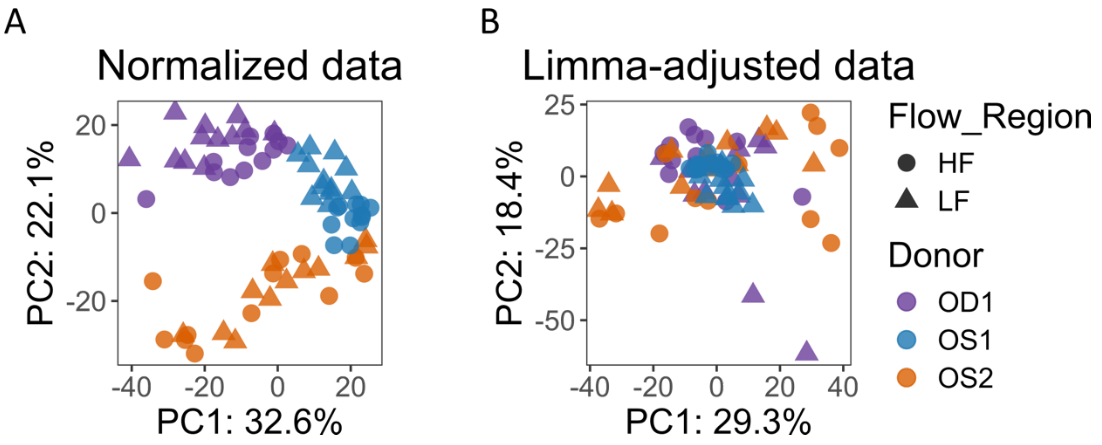


**Supplemental Figure 1: Principal Components Analysis before and after correcting for inter-donor correlation**. **A)** Principal Components Analysis on data after both Q3 and variance stabilizing normalization. The first two components (PC1 and PC2) are shown on the x and y axes. Each point represents a single ROI. The samples cluster by donor, shown by the colors, and not by flow region, shown by shape. **B)** PC1 and PC2 after the counts were adjusted for inter-donor correlation using limma in R. The ROIs no longer cluster by donor.

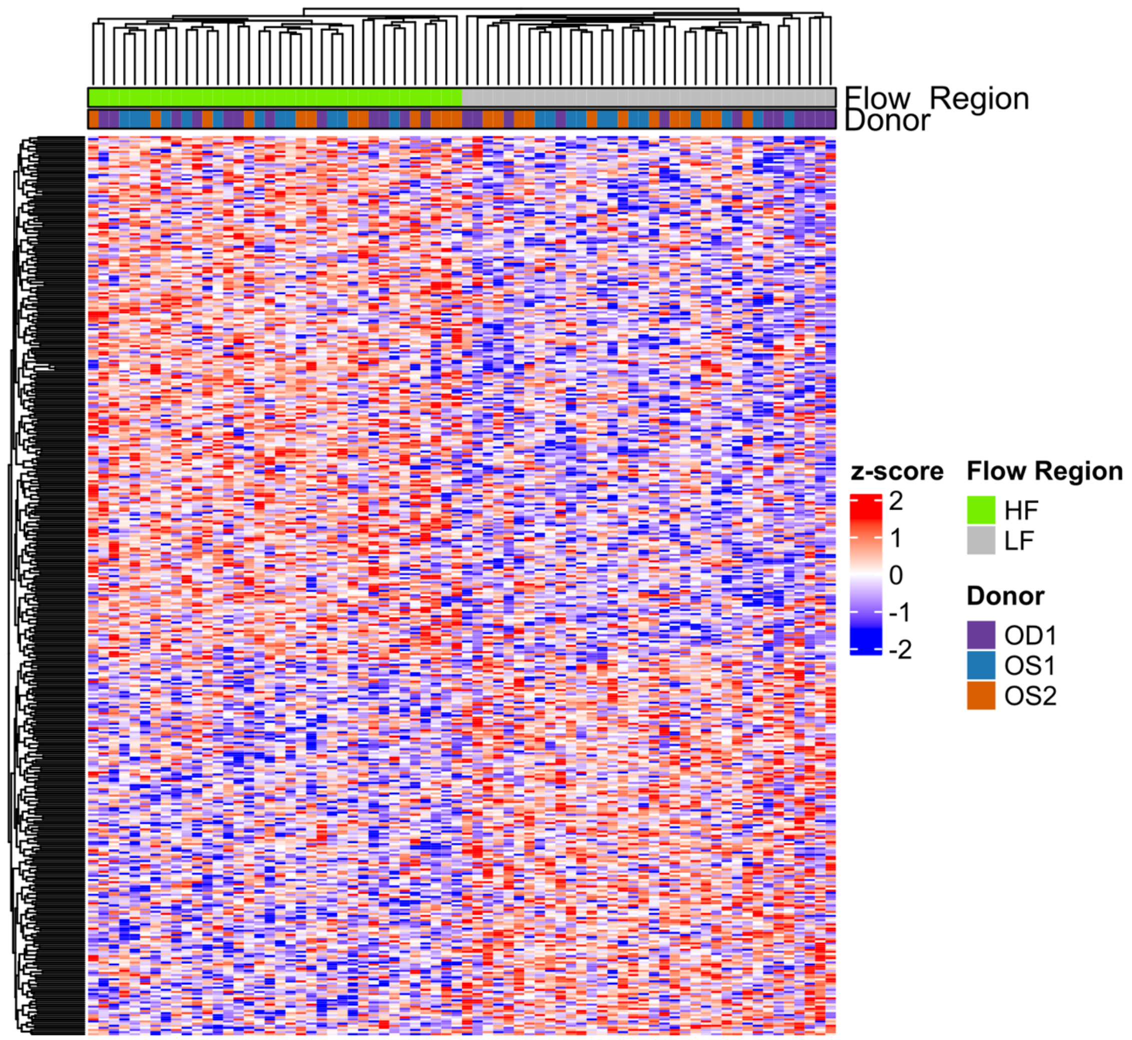


***Supplemental Figure 2: Heatmap of Human Differentially Expressed Genes.*** *Heatmap with hierarchical clustering of z-scored, normalized count data from all 426 differentially expressed genes. 249 genes are upregulated in HF, while 177 genes are upregulated in LF. There was no clustering by donor within up- and down- regulated genes.*

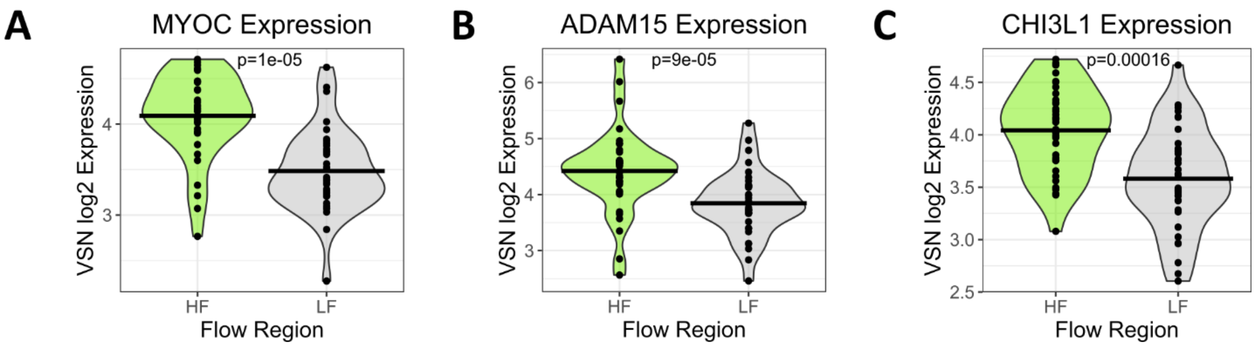


***Supplemental Figure 3: Top Differentially Expressed Genes in Human TM.*** *Violin plots comparing normalized log-transformed expression of myocilin, ADAM15, and CHI3L1 in HF vs. LF regions. Each point represents a single ROI.*

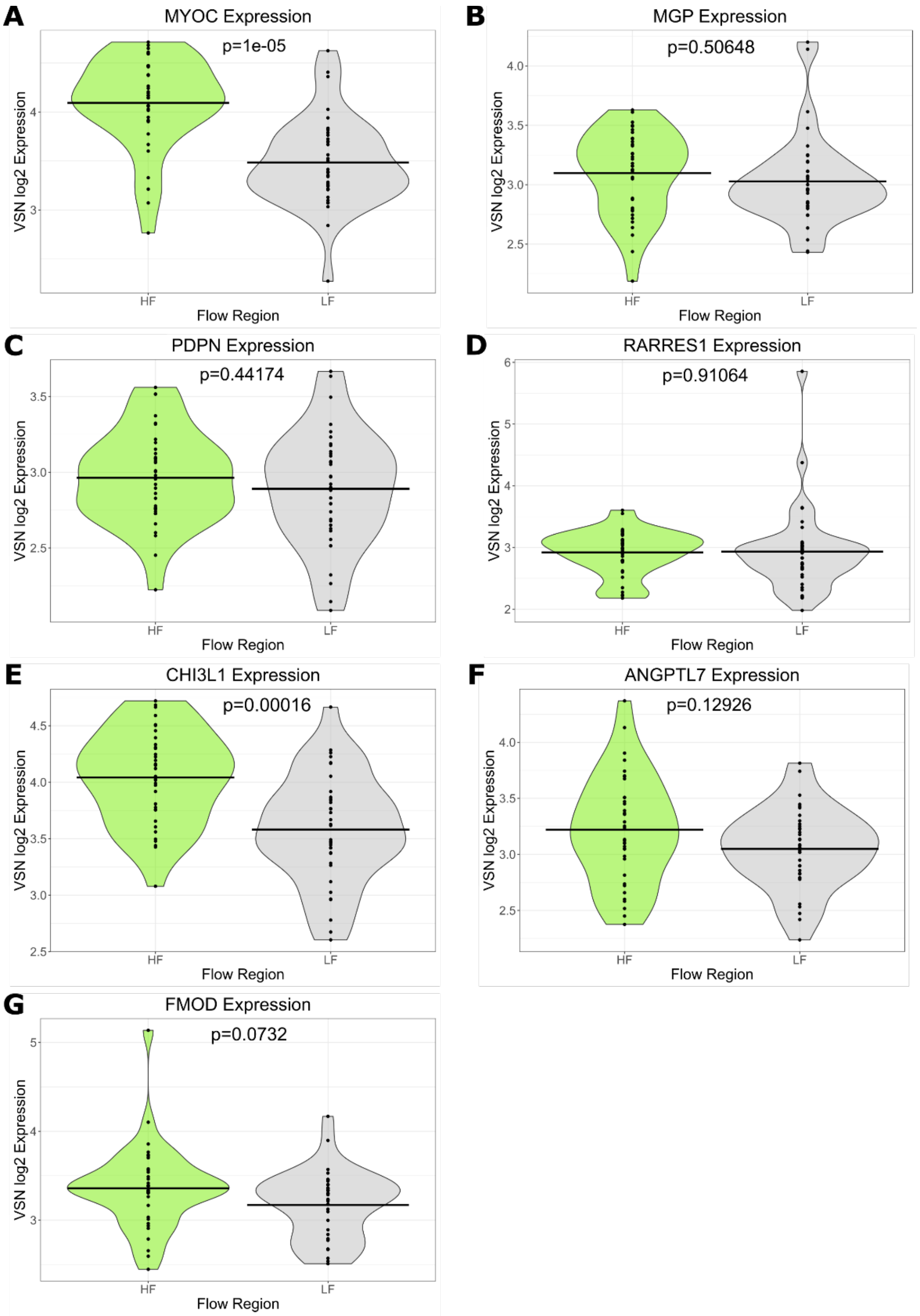


***Supplemental Figure 4:*** *Expression of TM cell markers in HF and LF regions.* ***A-D)*** *Canonical TM cell markers MYOC, MGP, PDPN, and RARRES1 expression levels in HF vs. LF regions. All four markers were expressed above the limit of quantification (LOQ) in our dataset.* ***E-G)*** *JCT cell markers CHI3L1, ANGPTL7, and FMOD expression levels in HF vs. LF regions. NELL2, the other JCT cell marker, was not detected above LOQ; however, the expression of 3 out of 4 JCT markers indicates that our data likely encapsulated JCT cells within the TM. Of*

*note, the main Beam A and Beam B-specific TM cell markers, FABP4 and TMEFF2, were not detected above the LOQ in our dataset.*

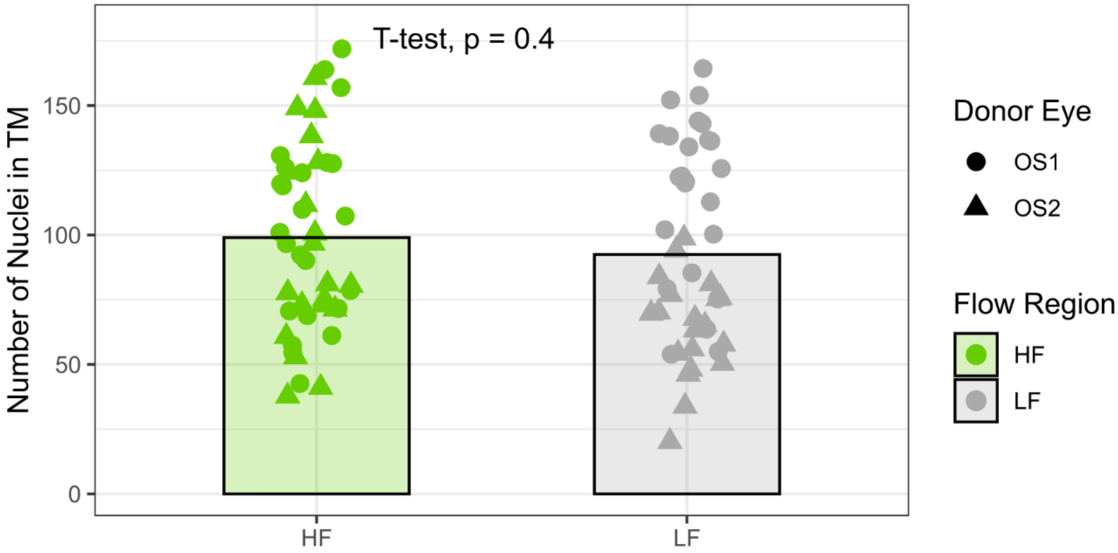


***Supplemental Figure 5: Nuclei counts in HF and LF Human TM.*** *No significant difference in nuclei counts was observed between sections from HF and LF regions (t-test, p = 0.42). Data includes 48 sections (24 HF, 24 LF) from Donor 1 OS eye and 40 sections (20 HF, 20 LF) from Donor 2 OS eye.*

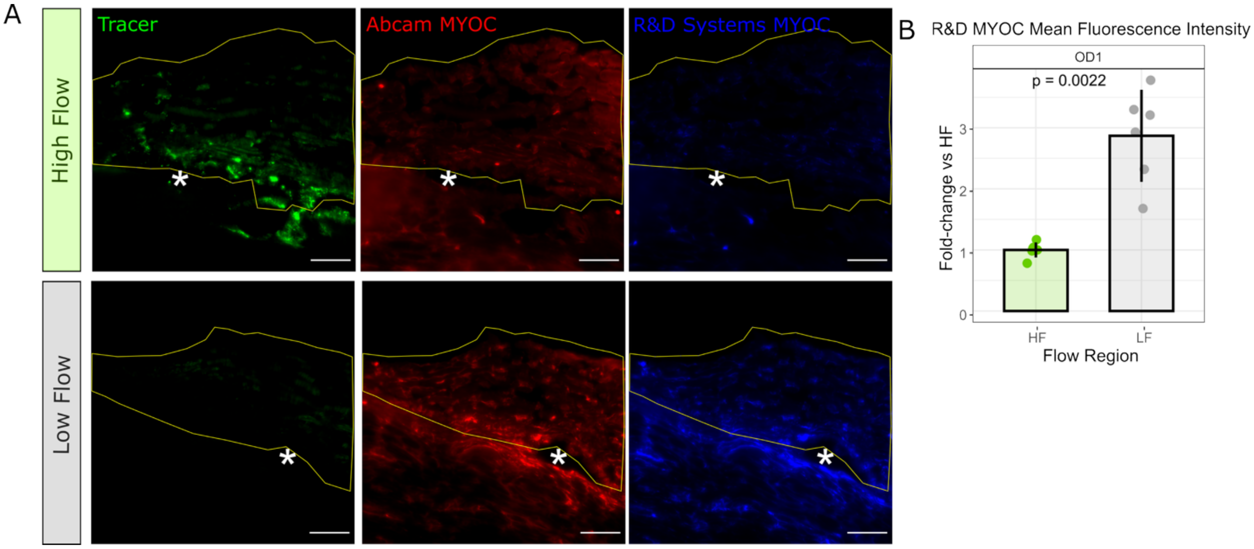


***Supplemental Figure 6: Myocilin Immunofluorescence Labeling with both R&D Systems and Abcam Primary Antibodies. A)*** *Representative HF and LF TM sections co-labeled with antibodies against myocilin from two sources: Abcam (red, as shown in Figure 3 of the main text) and R&D Systems (blue). Labeling was consistent between antibodies, with greater labeling present in LF TM and SC inner and outer wall. An asterisk denotes the SC lumen, the yellow outline shows the TM, and scale bars = 50 $\mu$m.* ***B)*** *Quantification of myocilin*

*labeling with the R&D Systems primary antibody in 6 HF and 6 LF sections from the OD eye of donor 1. Myocilin labeling was 2.87-fold greater in LF regions compared to HF regions ($p < 0.01$). p-values are obtained from Wilcoxon signed-rank tests, and bars represent the mean ± SD.*

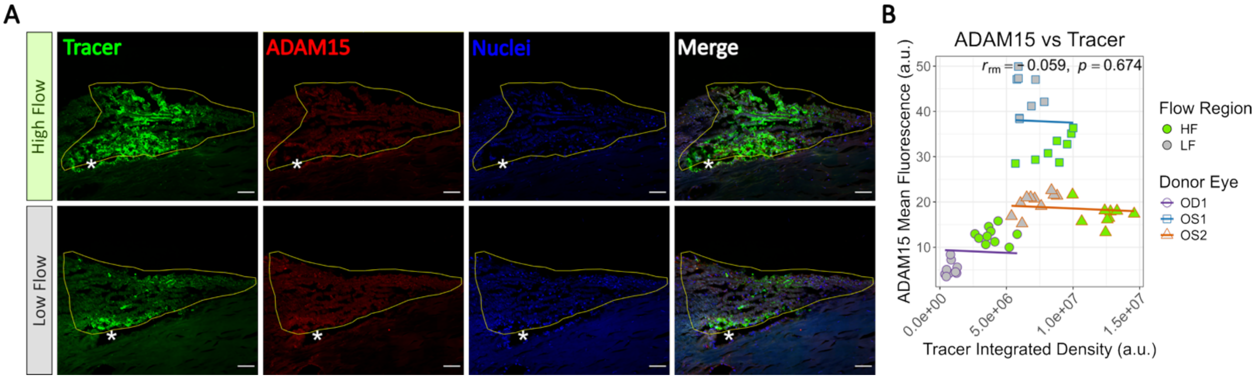


***Supplemental Figure 7: Immunofluorescent Staining of ADAM15 in Human HF and LF TM. A)*** *Sagittal sections of human TM from representative HF and LF regions of the Donor 2 OS eye labeled with ADAM15 (shown in red). The TM is outlined in yellow, and an asterisk (*) indicates the SC lumen. Scale bar = 50 μm. B) Repeated-measures correlation plot showing association between ADAM15 labeling vs. tracer intensity in the TM for all three donor eyes (OD1: n=10 HF and 9 LF sections, OS1: n=9 HF and 8 LF sections, OS2: n=10 HF and 10 LF sections). Points represent individual sections, and parallel lines indicate donor eye-specific regression fits ($r_{rm}$ represents the correlation coefficient; p-value represents repeated measures correlation significance). While some eyes individually showed differences in ADAM15 between segmental flow regions, no significant correlation was observed across all eyes.*

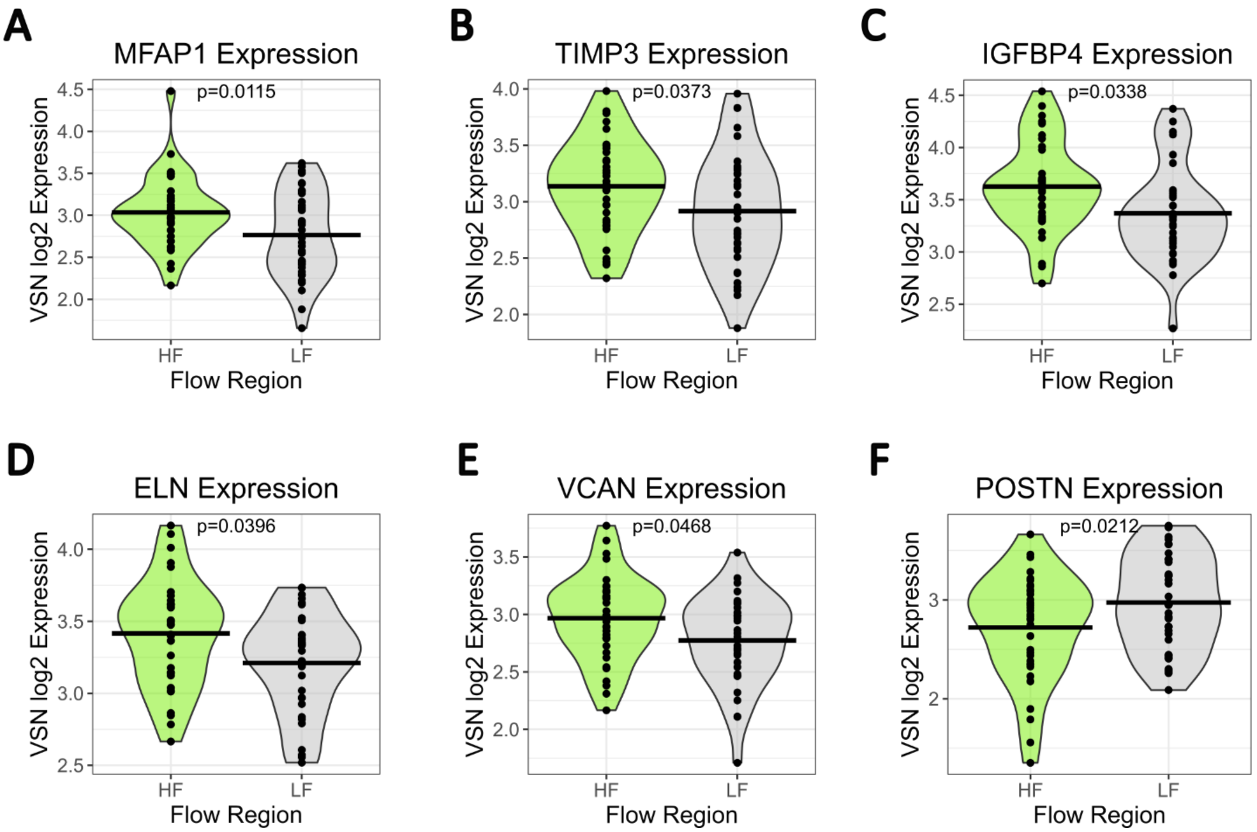


***Supplemental Figure 8: Differentially Expressed Matrix-Related Genes in Human TM.*** *Violin plots of normalized (VSN) log2-transformed expression values of differentially expressed ECM-related genes from HF and LF ROIs. MFAP1, TIMP3, IGFBP4, ELN, and VCAN were all upregulated in HF regions, while POSTN showed greater expression in LF regions. Each point represents a single ROI.*

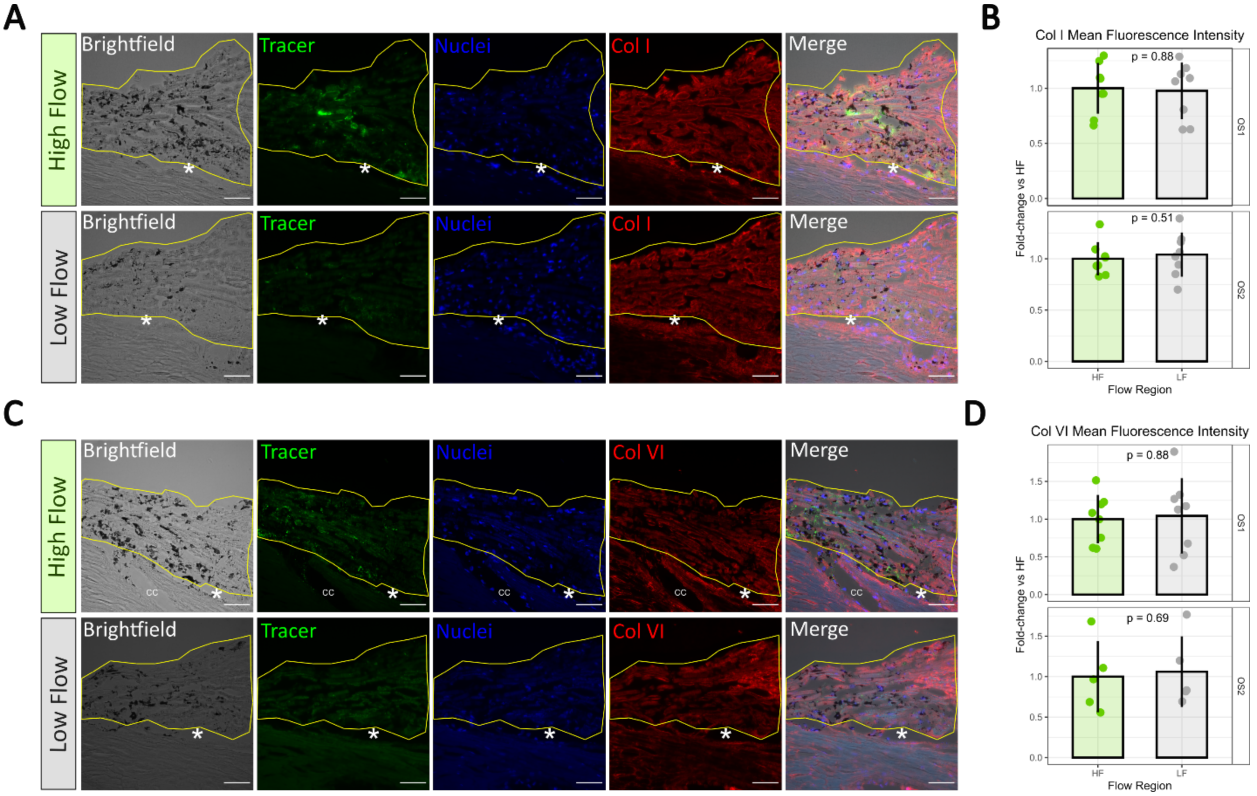


***Supplemental Figure 9: Collagens I and VI labeling is similar in HF and LF human TM.*** ***A)*** *Labeling of collagen I in representative HF and LF sections of human TM (Donor 1, OS eye).* ***B)*** *Quantification of collagen I mean fluorescence intensity in the TM of 8 HF and 8 LF sections from Donor 1 (OS eye), and 8 HF and 8 LF sections from Donor 2 (OS eye). No significant difference was found between HF vs. LF in either donor.* ***C)*** *Labeling of collagen VI in representative HF and LF sections of human TM (Donor 2, OS eye).* ***D)*** *Quantification of collagen VI mean fluorescence intensity in the TM of 8 HF and 8 LF sections from Donor 1 (OS eye), and 5 HF and 5 LF sections from Donor 2 (OS eye). No significant difference was found between HF vs. LF in either donor. P-values represent Wilcoxon tests, and bars represent the mean ± SD. Asterisks (*) denote the SC lumen and the yellow line outlines the TM (including SC inner wall). CC = collector channel. Scale bars = 50 µm.*

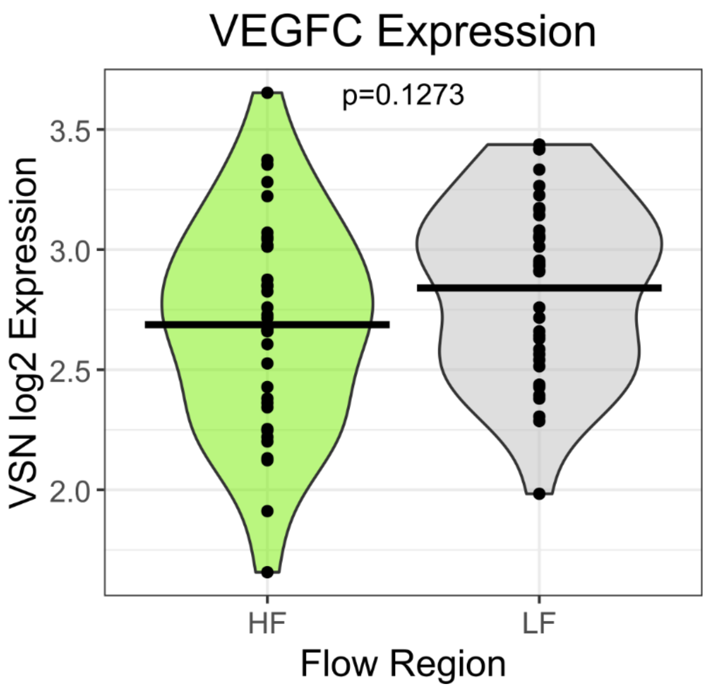


***Supplemental Figure 10: VEGFC is not differentially expressed between HF vs. LF regions of human TM.*** *Violin plot of normalized (VSN) log2-transformed expression values for VEGFC, with the p-value from the linear model used for differential expression analysis. Despite the differences observed in downstream VEGFA targets and VEGFB expression between HF and LF regions, we did not observe significant differential expression of VEGFC.*

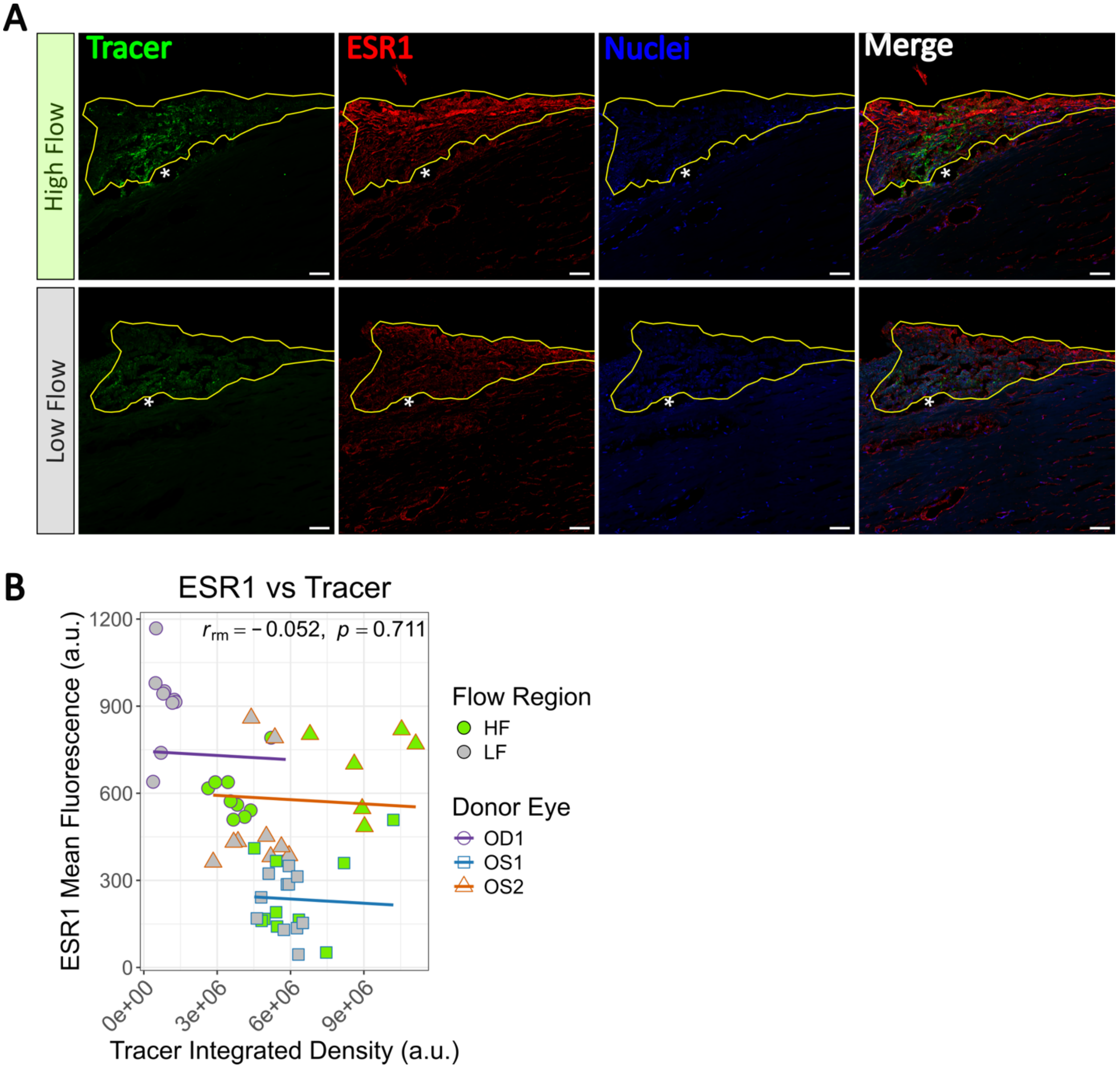


***Supplemental Figure 11: Immunofluorescent staining of ESR1 in human HF and LF TM. A)*** *Human TM sagittal sections (Donor 2, OS eye) from representative HF and LF regions labeled for ESR1. The TM is outlined in yellow, and the asterisk (*) denotes the SC lumen. Scale bar = 50 µm.* ***B)*** *Repeated-measures correlation plot showing association between ESR1 labeling vs. tracer intensity in the TM across all three donor eyes (OD1: n=10 HF and 9 LF sections, OS1: n=10 HF and 11 LF sections, OS2: n=6 HF and 9 LF sections). Points represent individual sections, and parallel lines indicate donor eye-specific regression fits with matched slopes ($r_{rm}$ represents the correlation coefficient; p-value represents repeated measures correlation significance). While some eyes individually showed differences in ESR1 between segmental flow regions, no significant correlation was observed across all eyes.*

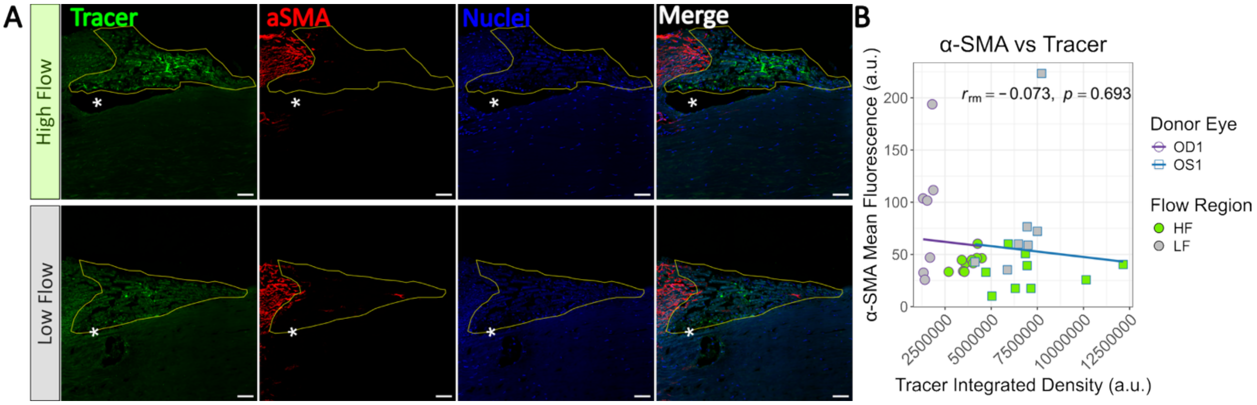


***Supplemental Figure 12: Immunofluorescent staining of $\alpha$-SMA in human HF and LF TM. A)*** *Human sagittal sections (Donor 1, OS eye) from representative HF and LF regions labeled for $\alpha$-SMA. The TM is outlined in yellow, and the asterisk (*) denotes the SC lumen. Scale bar = 50 µm.* ***B)*** *Repeated-measures correlation plot showing association between $\alpha$-SMA labeling vs. tracer intensity in the TM across both Donor 1 OS and Donor 1 OD eyes (n=10 HF and 8 LF sections per eye). Points represent individual sections, and parallel lines indicate donor eye-specific regression fits with matched slopes ($r_{rm}$ represents the correlation coefficient; p-value represents repeated measures correlation significance). In general, LF sections appeared to have slightly more $\alpha$-SMA compared to HF sections; however, HF sections were highly variably in both tracer and $\alpha$-SMA labeling. Overall, no significant correlation was observed between $\alpha$-SMA and tracer intensity.*

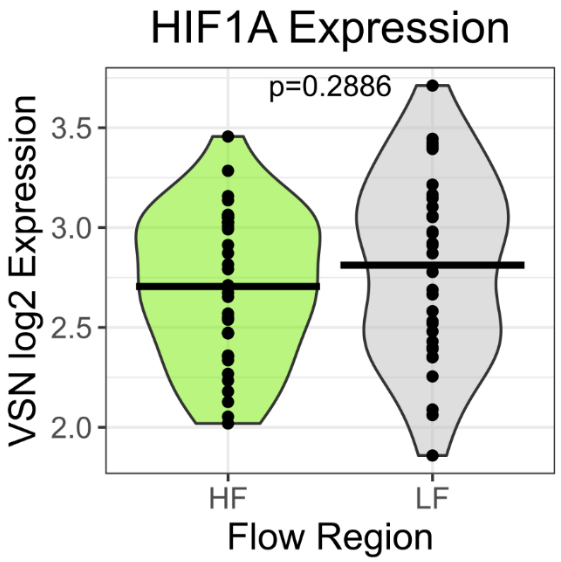


***Supplemental Figure 13: HIF1A is not differentially expressed in HF vs. LF regions of human TM.*** *Violin plot of normalized (VSN) log2-transformed expression values for HIF1A, with the p-value from the linear model used for differential expression analysis. Despite the differences observed in downstream HIF1A targets and hypoxia-related gene sets between HF and LF regions, we did not observe significant differential expression of HIF1A, which is considered a master regulator and key biomarker of hypoxia.*

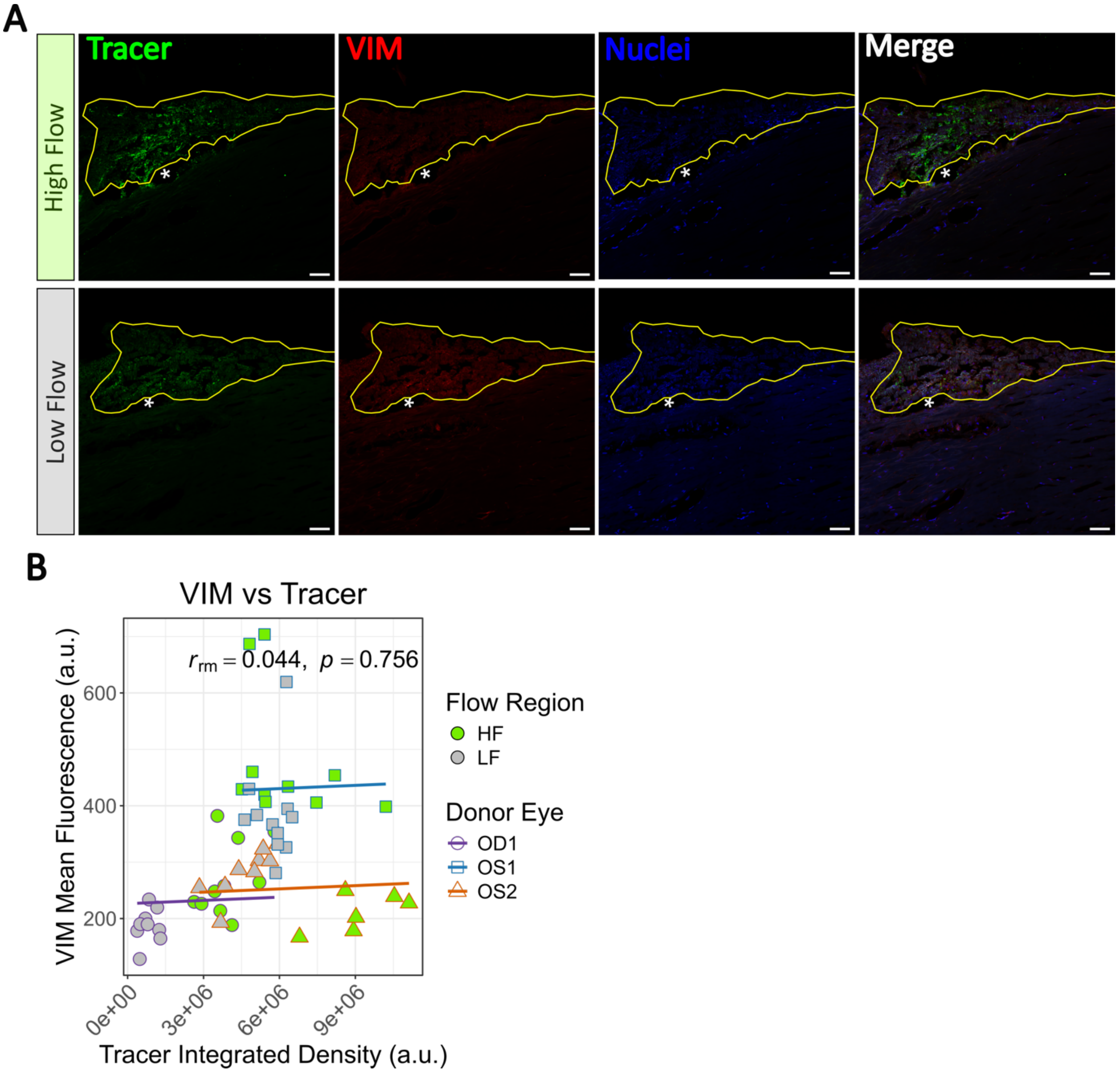


***Supplemental Figure 14: Immunofluorescent staining of Vimentin in Human HF and LF TM. A)*** *Human TM sagittal sections (Donor 2, OS eye) from representative HF and LF regions labeled for VIM. The TM is outlined in yellow, and the asterisk (*) denotes the SC lumen. Scale bar = 50 µm.* ***B)*** *Repeated-measures correlation plot showing association between VIM labeling vs. tracer intensity in the TM across all three donor eyes (OD1: n=10 HF and 9 LF sections, OS1: n=10 HF and 11 LF sections, OS2: n=6 HF and 9 LF sections). Points represent individual sections, and parallel lines indicate donor eye-specific regression fits with matched slopes ($r_{rm}$ represents the correlation coefficient; p-value represents repeated measures correlation significance). While some eyes individually showed differences in VIM between segmental flow regions, no significant correlation was observed across all eyes*
